# D1R-specific modulation of ACC mitigates chronic neuropathic pain

**DOI:** 10.64898/2025.12.31.697162

**Authors:** Yatika Chaudhary, Srividya Subash, Rajit Roy, Harikrishna Nair, Anant Jain, Arnab Barik

**Author notes:** Equal contribution (Yatika Chaudhary, Srividya Subash).

## Abstract

Maladaptive plasticity within central pain circuits is a defining feature of chronic neuropathic pain, yet the mechanisms governing these changes remain unclear. This study investigates the role of the mesocortical dopaminergic pathway in the neuropathic pain-induced hyperexcitability of dopamine D1 receptor-expressing anterior cingulate cortex (ACC^D1R^) neurons. We find that infusing D1R agonists in the ACC reverses the mechanical hypersensitivity and negative affective-motivational affect in mice caused by spared-nerve injury (SNI). Although ACC pyramidal neurons are widely presumed to be the principal targets of D1R signaling, ex vivo recordings reveal that D1R agonists suppress the excitability of D1R-expressing pyramidal neurons while enhancing the excitability of D1R-expressing interneurons. Consistently, gene-expression analyses show that D1R-expression is distributed across both excitatory and inhibitory ACC neurons. Furthermore, we demonstrate that the excitability of genetically labeled D1R pyramidal neurons is enhanced by SNI, whereas in interneurons, it is reduced. Chemogenetic manipulation demonstrates that activation of D1R neurons produces robust analgesic and anxiolytic effects, whereas inhibition worsens pain- and anxiety-related behaviours, indicating that inhibitory D1R neurons dominate population-level output. Circuit tracing further delineates diverse afferent and efferent connections linking ACC^D1R^ neurons to sensory and affective pain pathways. Together, these results identify dopaminergic regulation of ACC^D1R^ neurons as a critical determinant of cortical dysfunction in chronic pain.

## Introduction

Chronic neuropathic pain is a prevalent and debilitating condition affecting more than 5% of adults, yet it remains largely refractory to current therapies. One of the primary reasons for the failure of the development of successful treatment paradigms is the incomplete understanding of the neuronal pathophysiology that drives the hallmark symptoms: mechanical and thermal hypersensitivity, as well as stress and anxiety. Over the past few decades, research in animal models of neuropathic pain, such as spared nerve injury (SNI), diabetic neuropathy, and CIPN[9,14,24,61], has yielded substantial insight into the peripheral pathological processes; however, the alterations in central circuitries that sustain neuropathic pain remain elusive.

Imaging studies in human subjects have demonstrated that the ACC, an integral part of the medial prefrontal cortex, is spontaneously engaged in patients with neuropathic pain[21,29]. Mechanistic electrophysiological studies in mice and rats have shown that pyramidal neurons in the ACC are hyperexcitable under neuropathic conditions[57,75]. The potentiated ACC pyramidal neurons are attributed to the mechanical allodynia and negative emotional effects of chronic pain. The mechanism by which elevated activity in the ACC is induced and maintained in neuropathic conditions is a topic of active investigation. The neuromodulation provided by inputs from dopaminergic, serotonergic, and noradrenergic neurons in the midbrain and brainstem has been shown to influence the firing properties of ACC neurons[15,55]. Importantly, chronic pain results in a hypodopaminergic tone, negatively affecting motivated behaviours[62]. While L-DOPA administration rescued mechanical and thermal allodynia in neuropathic mice by indirectly reversing hyperexcitability in ACC pyramidal neurons through control of HCN-channel function[33]. Both dopamine receptors, D1R and D2R, are known to be expressed in the ACC; however, the D1R is critical in regulating the altered excitability in pyramidal neurons due to neuropathic pain[16,34,43]. It is unclear whether D1R is expressed in ACC interneurons and is subject to dopaminergic modulation. Notably, it was found that in the prefrontal cortex, D1R is expressed in Vip, PV, and Sst-expressing interneurons, in addition to pyramidal cells[2,39]. Thus, the excitability of ACC pyramidal and interneurons may be modulated by midbrain dopamine in neuropathic pain through D1R signaling.

Here, we have explored the roles of D1R receptor and D1R-expressing ACC neurons in mechanical allodynia and affective-motivational disorders resulting from peripheral neuropathy in mice with SNI. We tested the effects of the infusion of D1R agonists on the peripheral hypersensitivity and anxiety levels. Through ex vivo recordings, we tested the effects of D1R agonists on the excitability of excitatory and inhibitory ACC neurons. We asked if D1R modulation through pharmacologic antagonists and agonists altered field-LTP induction in the ACC. Furthermore, Cre recombinase expressed under the D1R promoter enabled us to express chemogenetic actuators and neuronal activity suppressors in the relevant ACC neurons, allowing us to test their role in the effects of SNI on affective-motivational behaviours and mechanical allodynia.

## Results

### In-vivo pharmacological activation of D1R receptor in ACC rescues SNI-induced hypersensitivity

Previous reports indicate that the ACC pyramidal neurons are under tonic dopaminergic modulation[33,34]. This modulation is inhibitory in nature and is facilitated by the D1R receptors expressed on the ACC pyramidal neurons [7]. However, the consequences of modulating D1R signaling in vivo on the mechanical allodynia and negative affective-motivational aspects of neuropathic pain remain unresolved. To test this, we utilized the spared nerve injury (SNI) model in mice to induce chronic neuropathic pain[8,17,48]. We tested the mechanical allodynia in the neuropathic mice with Von Frey filaments[19] and determined anxiety-like behaviors with the light-dark box test[7]. To facilitate D1R signaling in ACC neurons, we infused SKF 83822[44,47,64], a specific pharmacological D1R agonist, into the ACC through implanted cannula (Fig. 1A, B). Surprisingly, we found that compared to control saline-infused mice, SKF 83822 in wild-type CD1 mice increased mechanical thresholds and reversed mechanical allodynia post-SNI (Fig. 1C). Infusion of SKF 83822 in the ACC was anxiolytic in CD-1 mice. SNI was sufficient to increase anxiety, while infusion of SKF 83822 and not saline reduced the anxiogenic effects of SNI (Fig. 1D). Together, our data indicate that the activation of D1R signaling in ACC neurons, thereby mimicking receptor-specific dopamine-mediated modulation of ACC neurons, increases mechanical thresholds and decreases anxiety-like behaviors in both basal and neuropathic conditions.

**Figure 1:**
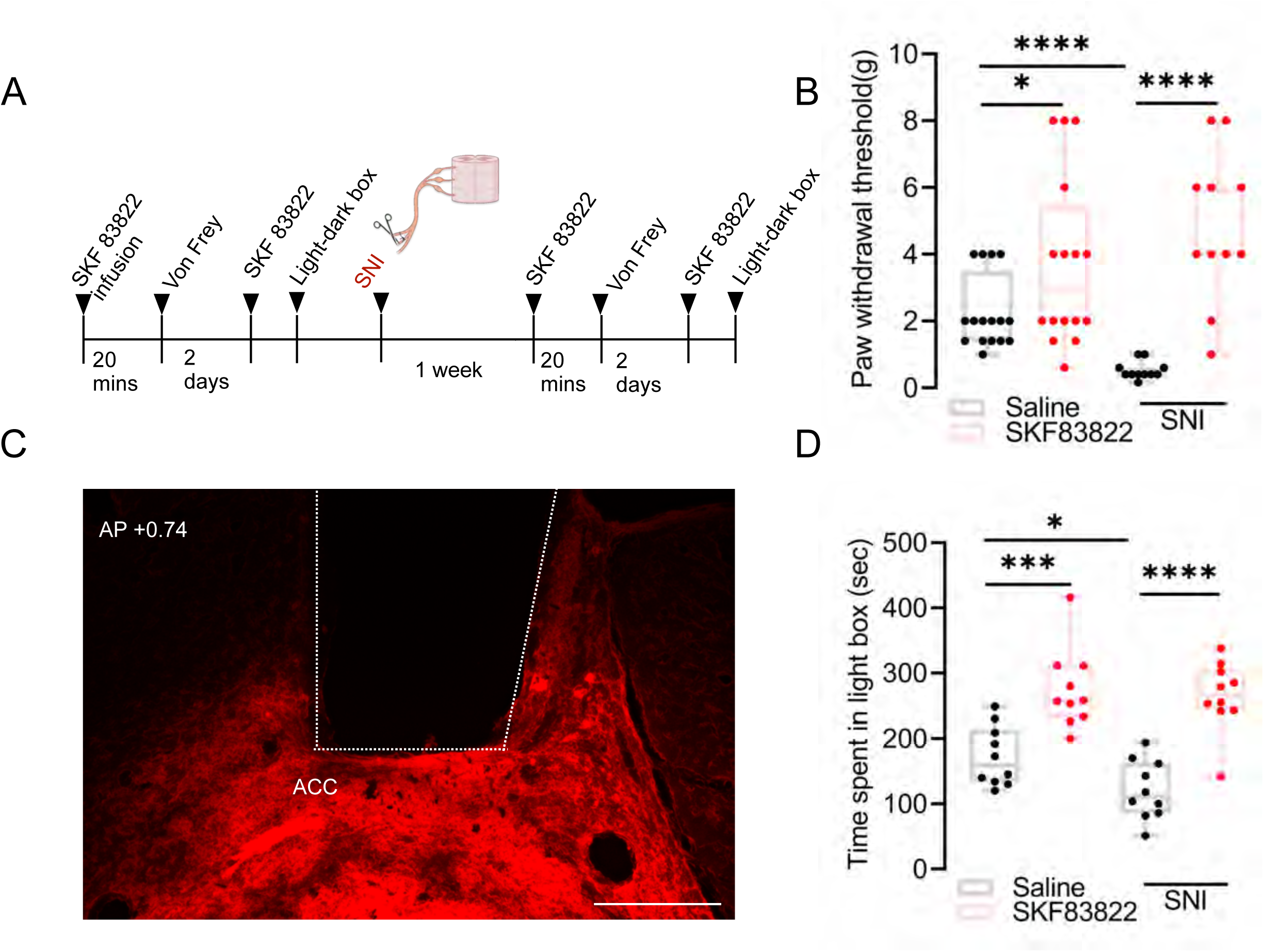
In-vivo pharmacological activation of D1R receptor in ACC rescues SNI-induced hypersensitivity. A) Schematic depicting the experimental timeline showing saline and SKF 83822 unilateral infusion in ACC before and after SNI through implanted cannulae, with corresponding behavioural assessments. B) Representative histological image of ACC showing the infusion site following SKF 83822 administration, visualised using CTB-555. C) Paw withdrawal threshold was assessed 20 minutes after saline or SKF 83822 infusion, comparing before (Unpaired t-test, 1.463±0.6879, *p=0.0418, n=16) and after (Unpaired t-test, 4.295±0.6764, ****p<0.0001, n=11) SNI. D) Time spent in the light box was assessed after 20 minutes of saline or SKF 83822 infusion, comparing before (Unpaired t-test, 102.4±24.08, ***p=0.0005, n=10) and after (Unpaired t-test, 1.44.3±22.20, ****p<0.0001, n=10) SNI.

### Synaptic plasticity in ACC is impaired following chronic pain and is dopamine-dependent

Chronic pain induces maladaptive plasticity in ACC neurons[5,42,76]. Peripheral inflammation, nerve injury, or digit amputation were shown to increase excitatory synaptic transmission in the deeper layers of ACC[37,72,74]. Moreover, theta-burst-induced LTP was occluded in mice with neuropathic pain[37]. Suggesting, ACC-mediated behavioral hypersensitivities due to neuropathic pain can be driven by altered plasticity in the ACC. In D1R agonist infusion behaviours we found D1R agonist has analgesic effect along with reduced anxiety (Fig. 1C,D), we hypothesized that the ACC plasticity could be dopamine dependent and is mediated by D1R. To test this hypothesis using acute slice electrophysiology, we measured the field excitatory postsynaptic potential (fEPSP) in ACC slices before and after SNI in CD1 mice (S1A). The magnitude of potentiation was estimated as the increase in fEPSP amplitude post-HFS. We found that the magnitude of potentiation in the SNI condition was significantly less than in the non-SNI condition (19% vs 64% respectively), suggesting that synaptic plasticity is impaired following chronic pain induction (S1B, C, D). Similarly, in the ACC^D1R^ neurons after paired-LTP induction, we did not observe significant potentiation in slices recovered from SNI mice (S2E-F). Furthermore, to test the role of dopamine in the regulation of synaptic plasticity in ACC, we used bath application of D1R antagonist (SCH23390) and D1R agonist (SKF83822) in non-SNI and SNI slices (Fig. 2A, E), respectively. We found that potentiation in the non-SNI condition was reduced with D1R antagonist (Fig. 2C, G), whereas D1R agonist application could not rescue the potentiation in the SNI condition (Fig. 2D, H). Therefore, we conclude that reduced dopamine activity in ACC can contribute to impaired synaptic plasticity in ACC, which occurs in chronic neuropathic pain.

**Figure 2:**
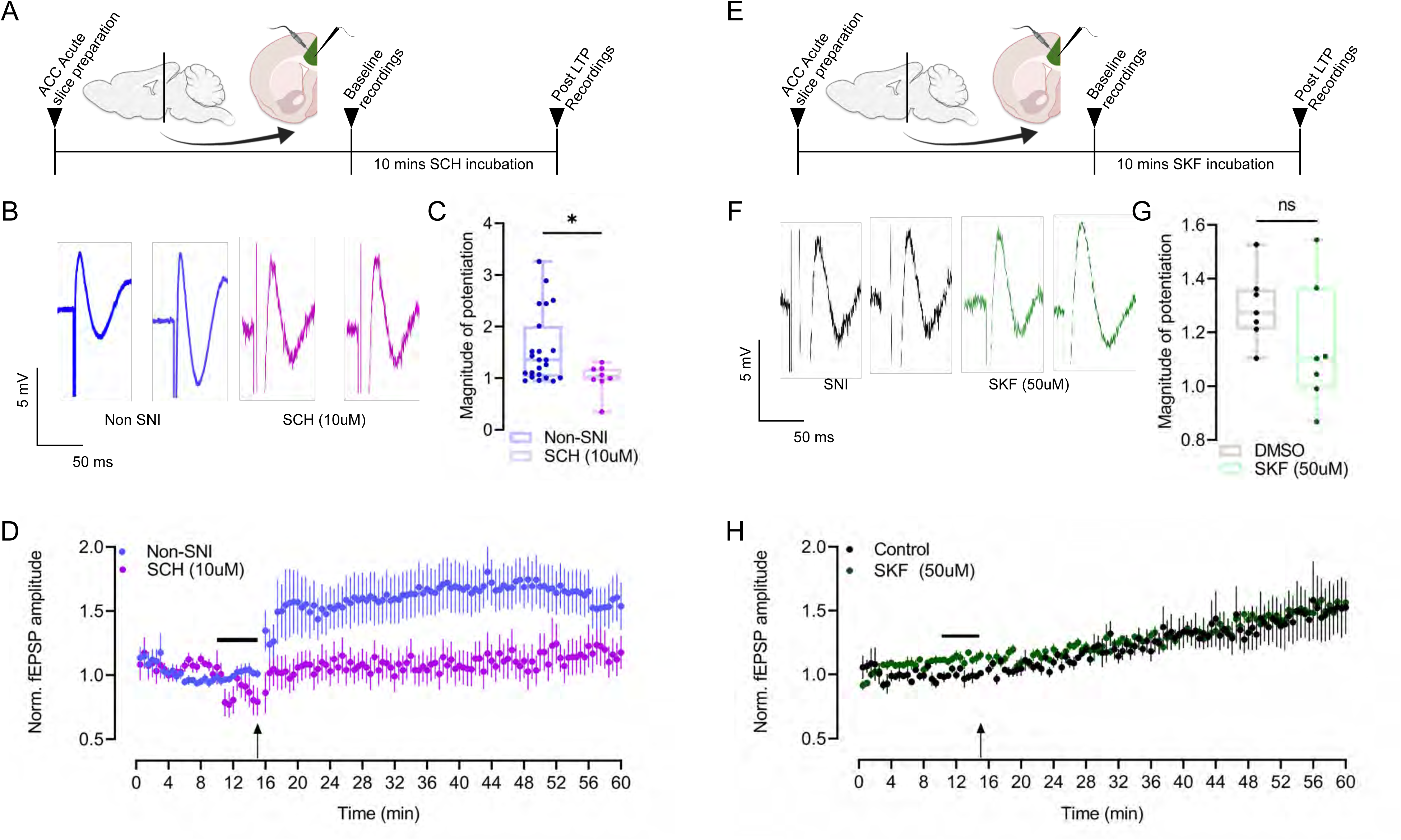
Synaptic plasticity in ACC is impaired following chronic pain and is dopamine-dependent. A) Schematics of field recordings from layer ⅚ in ACC with D1R antagonist application in basal condition. B) Representative traces showing fEPSP amplitude before and after LTP induction in basal conditions and with D1R antagonist. C) Comparison of magnitude of potentiation post 100 Hz HFS in basal condition (n=19) and with D1R antagonist (n=8), Unpaired t-test, −0.5616±0.2519, *p= 0.0337. D) Time course plot showing normalized fEPSPs amplitude with 15 mins baseline and followed by 45 mins recordings post 100 Hz HFS in basal condition and with D1R antagonist. E) Schematics of field recordings from layer ⅚ in ACC with D1R agonist application in SNI condition and with D1R agonist. DMSO mixed with aCSF was used as vehicle control in SNI slices. F) Representative traces showing fEPSP amplitude before and after LTP induction in SNI condition and with the D1R agonist. G) Comparison of magnitude of potentiation post 100 Hz HFS in SNI condition (n=12) and with D1R agonist (n=7), Unpaired t-test, p= 0.1157. H) Time course plot showing normalized fEPSPs amplitude with 15 mins baseline and followed by 45 mins recordings post 100 Hz HFS in the SNI condition and with D1R agonist.

### D1R-expressing ACC neurons are a heterogeneous population of excitatory and inhibitory neurons

While the existing literature is unclear regarding D1R expression across excitatory and inhibitory neuronal populations in the ACC, our functional data suggest that D1R-mediated modulation targets both regular-spiking and fast-spiking neurons. However, previous studies indicate that the D1R is expressed in both pyramidal and interneurons of the mPFC[54]. Here, we asked if, in a similar fashion, D1R is expressed both in the excitatory and inhibitory populations. To that end, we performed multiplex fluorescent in situ hybridization (RNAscope)[66] to probe for *drd1*,(gene encoding the D1R GPCR) and found that *drd1* expression was predominantly localised to the deeper cortical layers (V-VI) (Fig. 3A). The pyramidal ACC neurons are known to express *drd1*, whereas it is unclear if the interneurons express *drd1*. We found that *slc32a1*, the gene encoding Vgat, the vesicular inhibitory amino acid transporter, widely used as a marker for cortical inhibitory neurons, colocalizes with *drd1* (Fig. 3B). Furthermore, we genetically labeled the D1R-expressing neurons with eGFP, by stereotaxically co-injecting AAV-D1R-Cre (D1R promoter-driven Cre-recombinase)[3,20,52] with Cre-dependent AAV-DIO-eGFP (Fig. 3C). This strategy labeled the ACC^D1R^-expressing neurons with eGFP, and when we counterstained the brain sections with PV, a common marker for cortical interneurons, we found colocalization (Fig. 3C). We confirmed using RNAscope that the D1R-Cre indeed drove expression in the *drd1* neurons by probing for e*gfp* and *drd1* in brain slices recovered from mice which were infected with AAV-D1R-Cre and AAV-DIO-eGFP in the nucleus accumbens, where inhibitory neurons are known to express high levels of *drd1* (S2A). Thus, we can conclude from our gene expression analysis at both mRNA and protein levels that D1R receptors are present in both ACC pyramidal and interneurons.

**Figure 3:**
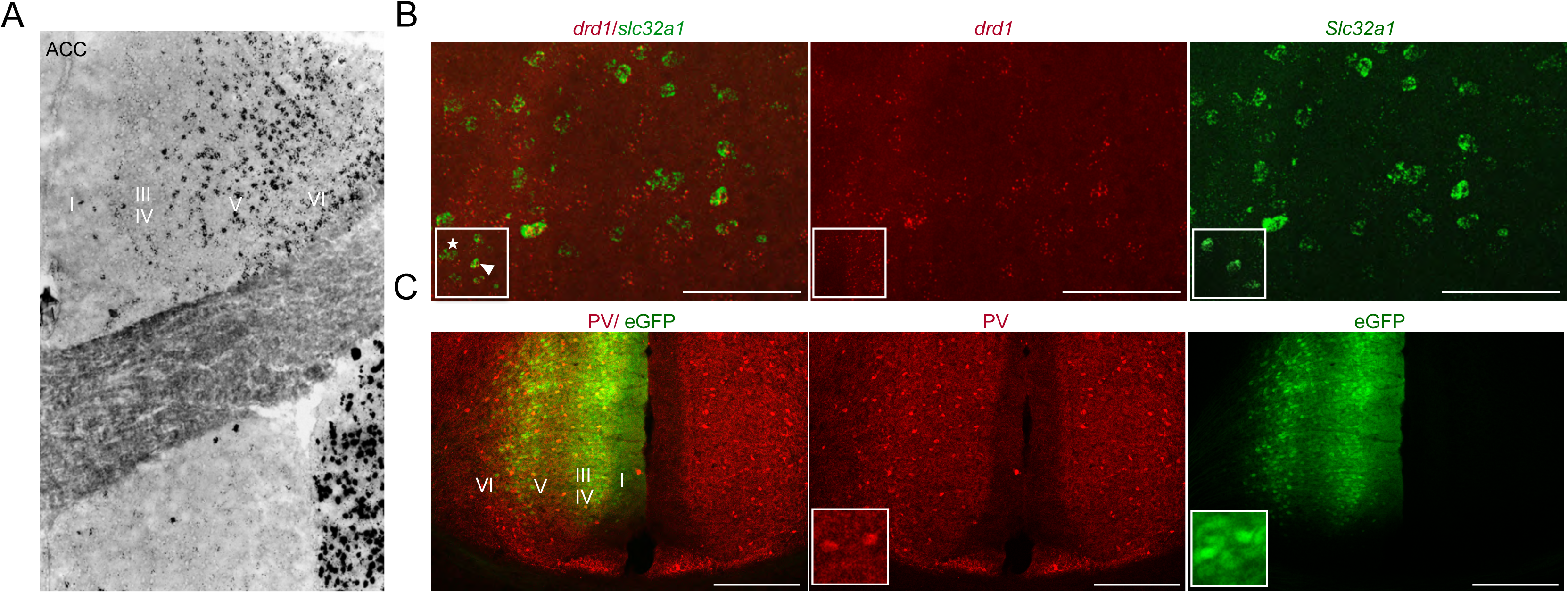
D1R-expressing ACC neurons are a heterogeneous population of excitatory and inhibitory neurons. A) Multiplex in-situ hybridisation showing drd1 expression profile along different cortical layers of ACC. B) Multiplex in-situ hybridisation showing co-expression of drd1(red) within slc32a1 (green) neurons in the ACC. C) PV staining in ACC sections from mice coinjected with AAV encoding D1R Cre and DIO eGFP. Yellow cells indicate co-expression of Paravalbumin and eGFP in ACC neurons.

### Nerve injury-induced potentiation of ACC neurons can be scaled down with D1R agonist in ex vivo recordings

To understand the mechanistic basis of SKF 83822’s effects on ACC neurons, we performed ex vivo whole-cell patch clamp electrophysiology recordings in unlabelled ACC neurons. Previous studies indicate that pyramidal neurons are rendered hyperexcitable due to disturbed D1R mediated signaling following SNI in rodents[6,12]. Therefore, we reasoned that D1R agonist application in the ACC could have a suppressive effect on the excitability of ACC pyramidal neurons. When we recorded from Layer ⅚ ACC neurons of CD-1 mice with SNI (Fig. 4A, B), we found that SKF 83822 had opposing effects on the excitability of regular and fast-spiking cells. Regular spiking neurons showed reduced firing, whereas fast spiking neurons showed increased firing at higher current injection steps (Fig. 4C, D, I, J). We recorded 79 ACC neurons (unlabelled and D1R labelled) comprising 67.0886% regular spiking and 32.911% fast spiking neurons (S3A, B). We segregated these two populations on the basis of number of action potentials, action potential spike half width, decay time and action potential amplitude. Neurons which showed action potentials above 13 at 200pA tends to be fast spiking (putatively interneuron), whereas neurons firing action potential within 13 are labelled as regular spiking (putatively excitatory neurons) (S3 C-F)[27,45].. The suppressive effects of SKF 83822 on the pyramidal neurons were expected; however, the effect on D1R-modulation on the ACC interneurons was surprising. The simultaneous bidirectional effects of SKF 83822 on the ACC pyramidal and interneurons may explain the results of the infusion experiments (Fig. 4C, D). Apart from the firing changes, passive membrane properties, such as rheobase and input resistance, remained unchanged in both regular spiking and fast spiking interneurons (Fig. 4G, M, H, N). Together, these observations revealed that dopamine modulates the ACC local circuitry by potentially regulating the excitability of both regular (excitatory) and fast-spiking (inhibitory) neurons.

**Figure 4.**
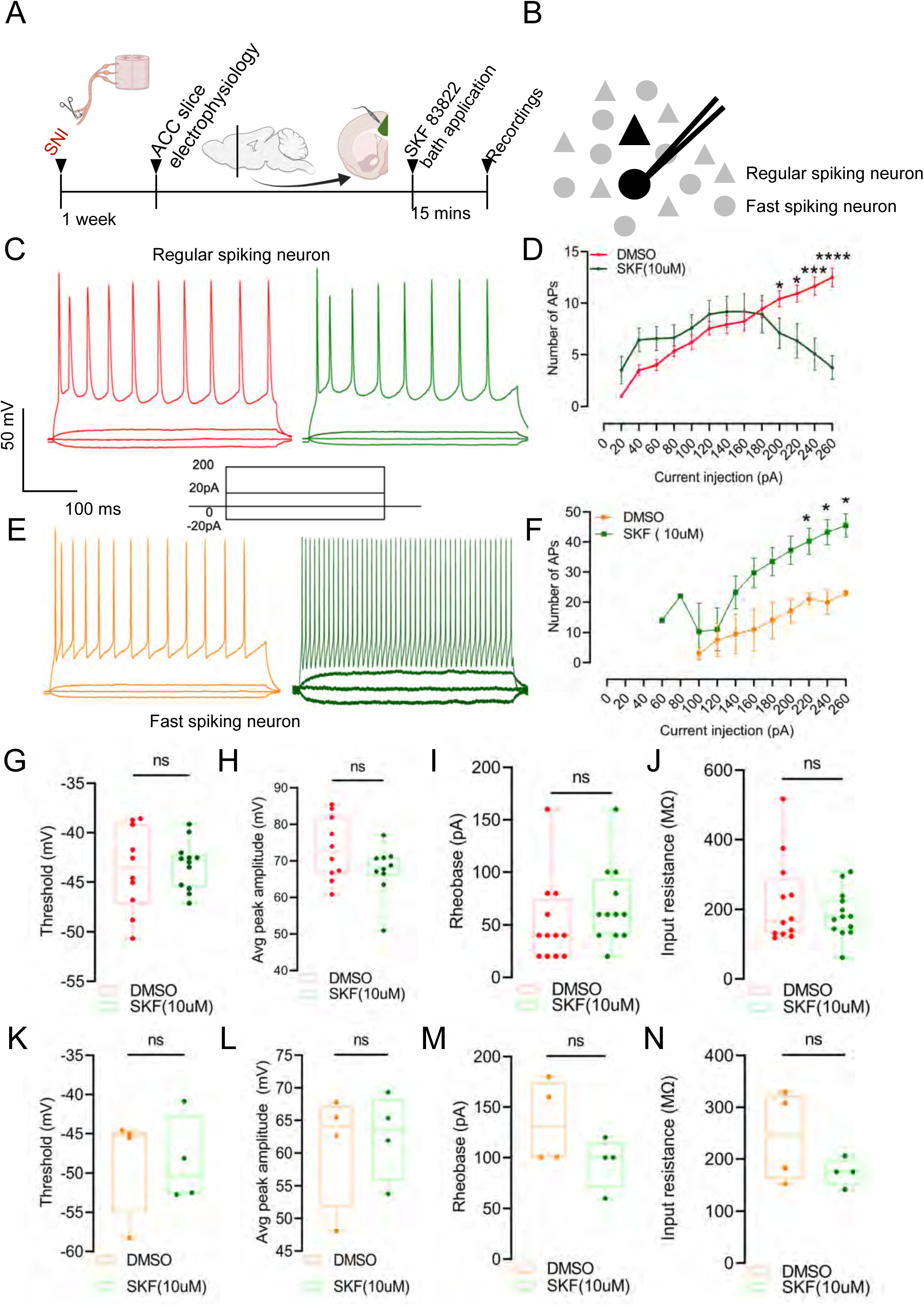
Nerve injury-induced potentiation of ACC neurons can be scaled down with D1R agonist in ex vivo recordings. A) Schematics showing whole cell patch clamp recordings from regular spiking and fast spiking neurons in ACC. B) Schematics showing paradigm of whole cell patch clamp recordings from ACC slices recovered from SNI mice. C) Sample action potential recordings from ACC regular spiking neurons (RSN) before and after SKF bath application at respective step currents (−20, 0, 20, 200 pA). D) Comparison of RSN action potential firing between DMSO and SKF (Unpaired t-test, 3.769±1.570, *p= 0.0244, 5.000±1.753, **p= 0.0088, 6.769±1.688, *** p= 0.0005, 8.615±1.472, ****p< 0.0001, n= 13). E) Sample action potential recordings from ACC fast spiking neurons (FSN) before and after SKF bath application at respective step currents (−20, 0, 20, 200 pA). F) Comparison of FSN action potential firing between DMSO (Unpaired t-test, 22.00±5.583, **p= 0.0076, 21.000±4.979, **p= 0.0056, 21.00±4.486, **p= 0.0034, 23.00±4.505, **p=0.0022, 22.75±3.955, **p=0.0012, n= 4). G) Comparison of RSN threshold between DMSO and SKF (Unpaired t-test, p= 0.9980, n= 10). H) Comparison of RSN average peak amplitude between DMSO and SKF(Unpaired t-test, p= 0.1712, n= 10). I) Comparison of RSN rheobase between DMSO and SKF (Unpaired t-test, p= 0.3071, n= 12). J) Comparison of RSN input resistance between DMSO (n= 12) and SKF (n= 13), Unpaired t-test, p= 0.3850. K) Comparison of FSN threshold between DMSO and SKF (n= 4), Unpaired t-test, p= 0.9637. L) Comparison of FSN average peak amplitude between DMSO and SKF (n= 10), Unpaired t-test, p= 0.7847. M) Comparison of FSN rheobase between DMSO and SKF (n= 12), Unpaired t-test, p= 0.1488. N) Comparison of FSN input resistance between DMSO (n= 12) and SKF (n= 13), Unpaired t-test, p= 0.1883.

### Nerve injury renders ACC^D1R^ excitatory neurons hyperexcitable, while dampening the excitability of inhibitory neurons

Nerve injury-induced hyperexcitability in the ACC pyramidal neurons has been reported via in vivo and ex vivo recordings in rodent models, and this excitability is modulated by D1R[12,33,34]. In our whole cell patch clamp electrophysiology recordings with D1R agonist, we found that the action potential firing of regular spiking (excitatory) and fast spiking (inhibitory) neurons were modulated differently (Fig. 2C, D, I, J). Based on these findings and the differential expression of *drd1*, we hypothesized that this hyperexcitability in the D1R-expressing ACC neurons in neuropathic conditions may result from reduced inhibition or a disturbance in the excitatory-inhibitory (E/I) balance. To test this hypothesis, we performed whole-cell patch-clamp electrophysiology on fluorescently labeled (eGFP) ACC^D1R^ neurons[3,20,52] in acute brain slices obtained from mice with and without spared nerve injury (Fig. 5A, B). Electrophysiological recordings from the eGFP-expressing ACC^D1R^ neurons revealed the diversity in the neuronal firing in control animals. SNI demonstrated contrasting effects on regular and fast-spiking neurons. While regular spiking neurons exhibited an increase in firing with higher current injections, fast spiking interneurons showed a decrease in firing in response to higher current injections (Fig. 5C, D, E, F). This increased excitability of regular spiking neurons was complemented with higher input resistance (Fig. 5J) and a hyperpolarizing shift in rheobase membrane potential (Fig. 5I). However, the input resistance and rheobase remained unchanged after SNI in the fast-spiking neuron (Fig. 5N, M). Furthermore, the action potential firing threshold and peak amplitude remained unchanged in both populations after spared nerve injury (Fig. 5G, K, H, L). Overall, SNI exhibited contrasting effects on the intrinsic excitability of regular spiking neurons versus fast-spiking neurons. These electrophysiology results suggest that both the hyperexcitability of D1R-expressing pyramidal neurons and the reduced firing of D1R-expressing inhibitory neurons contribute to SNI-induced mechanical allodynia and pain-associated negative emotional states.

**Figure 5.**
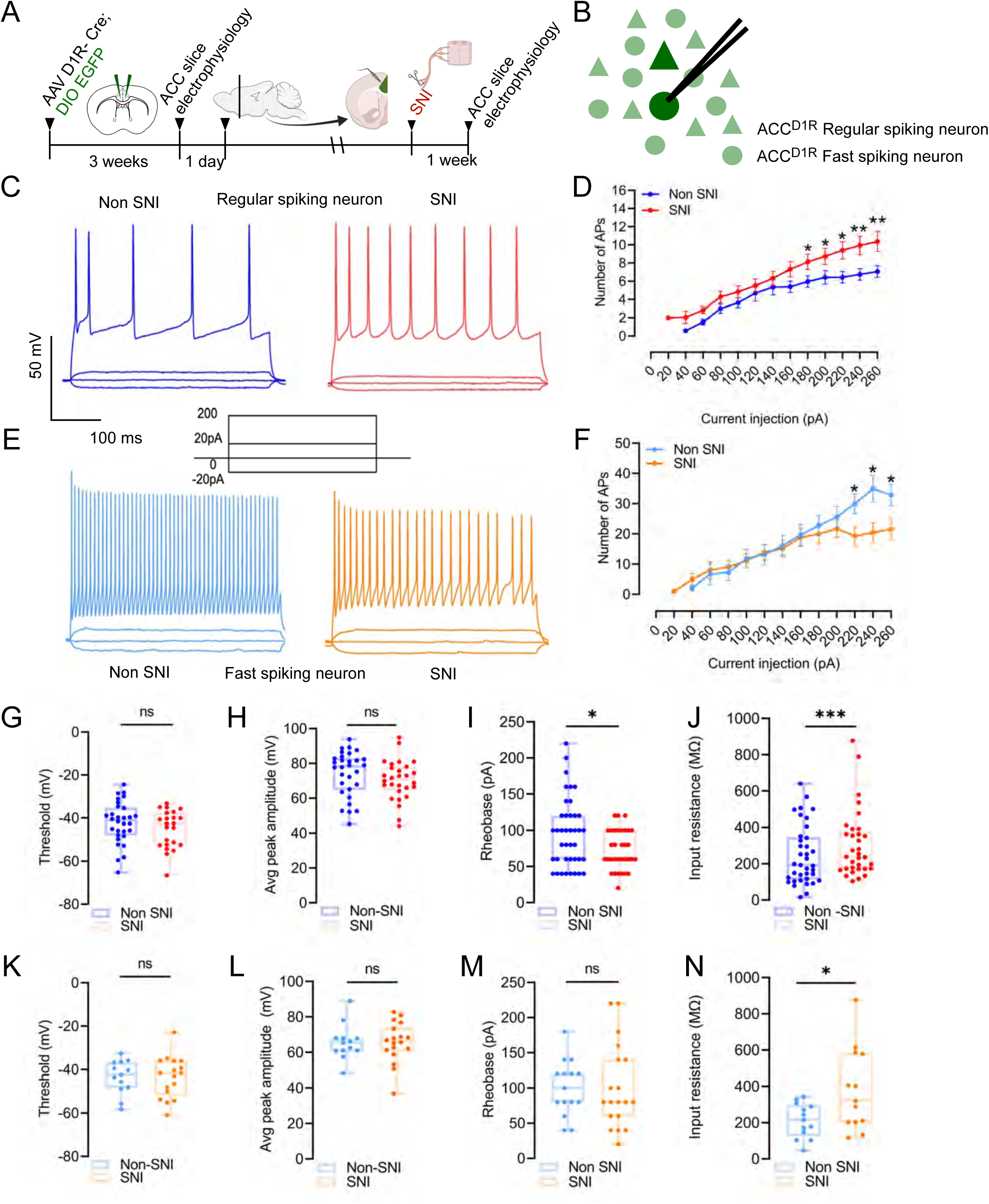
Nerve injury renders ACC^D1R^ excitatory neurons hyperexcitable, while dampening the excitability of inhibitory neurons. A) Schematics of whole cell patch clamp recordings from D1R-labelled ACC neurons in baseline conditions. B) Schematics of whole cell patch clamp recordings from D1R-labelled ACC neurons in SNI conditions. C) Sample action potential recordings from ACC regular spiking neurons (RSN) during baseline and after SNI at respective step currents (−20, 0, 20, 200 pA). D) Comparison of RSN action potential firing during baseline (n= 40), and after SNI (n= 33), Unpaired t-test, 2.164±1.024, *p= 0.0383, 2.311±1.1157, *p= 0.0497, 2.961±1.121, *p= 0.0102, 3.194±1.160, **p= 0.0075, 3.303±1.219, **p= 0.0084. E) Sample action potential recordings from ACC fast spiking neurons (FSN) during baseline and after SNI at respective step currents (−20, 0, 20, 200 pA). F) Comparison of FSN action potential firing during baseline (n= 15), and after SNI (n= 18), Unpaired t-test, −10.68±4.488, *p= 0.0236, −14.47±5.370, * p= 0.0113, −11.31±4.906, *p= 0.0280. G) Comparison of RSN threshold during baseline (n= 30), and after SNI (n= 23), Unpaired t-test, p= 0.2375. H) Comparison of RSN average peak amplitude during baseline (n= 30), and after SNI (n= 28), Unpaired t-test, p= 0.2403. I) Comparison of RSN rheobase during baseline (n= 40), and after SNI (n= 33), Unpaired t-test, −22.17±9.312, *p= 0.0200. J) Comparison of RSN input resistance during baseline (n= 37), and after SNI (n= 33), Unpaired t-test, −128.6±46.0 **p= 0.0067. K) Comparison of FSN threshold during baseline (n= 13), and after SNI (n= 17), Unpaired t-test, p= 0.9607. L) Comparison of FSN average peak amplitude during baseline (n= 13), and after SNI (n= 18), Unpaired t-test, p= 0.9879. M) Comparison of FSN rheobase during baseline (n= 15), and after SNI (n= 18), Unpaired t-test, p= 0.7422. N) Comparison of FSN input resistance during baseline (n= 13), and after SNI (n= 13), Unpaired t-test, 168.5±68.38, *p=0.0213.

### Activating ACC^D1R^ neurons alleviates SNI-induced mechanical allodynia

In the next set of experiments, we manipulated the activity of the ACC^D1R^ neurons. We reasoned that altering the activity of the ACC^D1R^ neurons in mice with neuropathic pain would inform us about their functional implications on mechanical allodynia. To that end, we injected AAVs encoding Cre-dependent hM3Dq-Cherry and D1R-Cre bilaterally in the ACC of CD-1 mice (Fig. 6A). This strategy enabled the chemogenetic activation of ACC^D1R^ neurons upon intraperitoneal (i.p.) administration of DCZ[40,50]. We confirmed hM3Dq expression by visualizing the tagged mCherry and successful chemogenetic stimulation by assaying for IEG cFos[25,59] expression upon DCZ-mediated neuronal activation (Fig. 6B). We utilized Von Frey filaments to provide graded punctate stimuli to the hind paws in mice to test for static mechanical threshold and the brush assay to test for dynamic mechanical pain threshold[10]. We found that upon chemogenetic activation of the ACC^D1R^ neurons, the Von Frey and brush assay thresholds remained unchanged; however, when mechanical allodynia was induced by performing SNI in mice, chemogenetic activation of the ACC^D1R^ neurons increased the thresholds, providing analgesia (Fig. 6C, D). In previous sections, we determined that the ACC^D1R^ neurons comprise a mixture of pyramidal and interneurons (Fig. 3), and that pyramidal neurons are rendered hyperexcitable by SNI (Fig. 5). Thus, we hypothesized that the allodynia caused by the ACC^D1R^ neurons is due to the dominant effect of stimulating the inhibitory interneurons, as stimulating only the pyramidal neurons would be expected to have a hyperalgesic effect. Indeed, when we chemogenetically activated the CaMKII-expressing ACC pyramidal neurons by stereotaxically delivering AAV-CaMKII-Cre[56] and AAV-DIO-hM3Dq in the ACC and administering DCZ i.p., we found that it was sufficient to cause mechanical hyperalgesia (Fig. 6E, F). Von Frey and Dynamic Brush thresholds were found to be lower in mice without SNI (Fig. 6G, H). After SNI, there were no observable effects of activating the ACC CaMKII neurons due to the already lowered mechanical thresholds in the Von Frey test, whereas in the Dynamic Brush test, the analgesic effects of targeted activation were observable (Fig. 6H). Whereas, when we chemogenetically stimulated the interneurons in the ACC by driving hM3Dq-mCherry expression (Fig. S4A, B) under the Vgat promoter, it caused analgesia under baseline and neuropathic conditions (Fig. S4C). Thus, stimulating interneurons is sufficient to increase mechanical thresholds. Importantly, next, when we stimulated the CaMKII+ve pyramidal and Vgat+ interneurons simultaneously with targeted chemogenetics, we observed analgesia under baseline conditions as well as in mice with SNI (Fig. S4D-F). Thus, when ACC pyramidal and interneurons are activated together, the effect of interneuron stimulation on mechanical thresholds can potentially override the roles played by pyramidal neurons.

**Figure 6.**
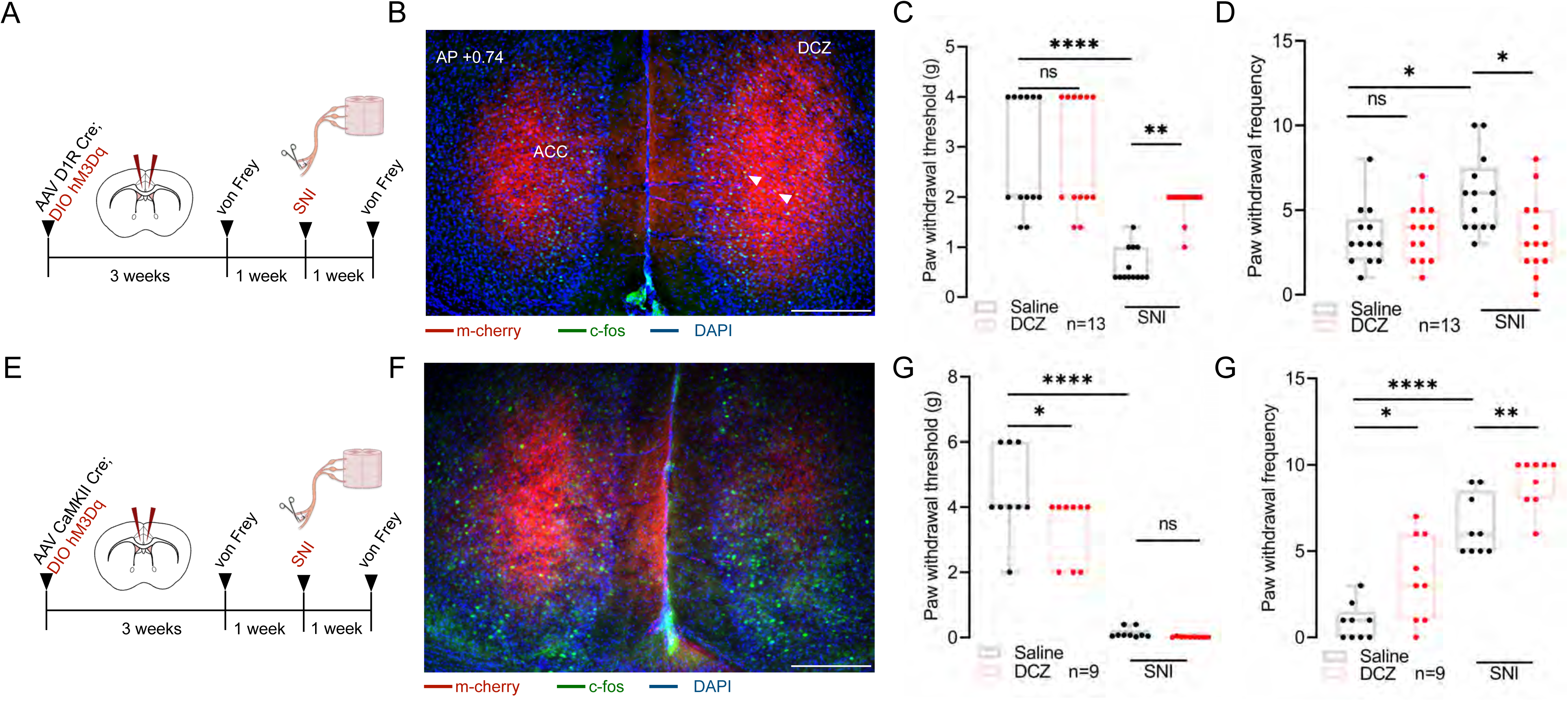
Activating ACC^D1R^ neurons alleviates SNI-induced mechanical allodynia. A) Schematic of experimental design. AAV encoding D1R^Cre^ and Cre-dependent excitatory DREADD hM3Dq was injected bilaterally into the ACC of CD1 mice. B) Fluorescence microscope image of AAV DIO hM3Dq bilateral expression in ACC D1R neurons. C) Paw withdrawal threshold pre (Tukey’s multiple comparison, p >0.9999) and post (Tukey’s multiple comparison, −1.246± 0.3305, *p=0.0024) SNI upon DCZ administration (n=13). D) Paw withdrawal frequency pre (Tukey’s multiple comparison, p =0.9974) and post (Tukey’s multiple comparison, 2.462± 0.7976, *p=0.0171) SNI upon DCZ administration (n=13). E) Schematic of experimental design. AAV encoding CAMKII Cre and Cre-dependent excitatory DREADD hM3Dq was injected bilaterally into the ACC of CD1 mice. F) Fluorescence microscope image of AAV DIO hM3Dq bilateral expression in ACC CAMKII neurons. G) Paw withdrawal threshold pre (Paired t-test, −1.111± 0.3514, *p=0.0133) and post (Paired t-test, p=0.0549) SNI upon DCZ administration (n=9). H) Paw withdrawal frequency pre (Paired t-test, 2.556± 0.5800, **p=0023) and post (Paired t-test, 2.556± 0.6479, ** p= 0.0043) SNI upon DCZ administration (n=9).

### Silencing ACC^D1R^ neurons is sufficient to induce mechanical allodynia

Since activation of the ACC^D1R^ neurons causes allodynia, we asked what would happen to the somatosensory mechanical thresholds if these neurons were inactivated. To that end, we expressed inward rectifying GFP-tagged Kir2.1[49,73] channels in a Cre-dependent manner under the D1R promoter bilaterally in the ACC neurons of CD-1 mice (Fig. 7A, B). As controls, we expressed GFP in the ACC neurons. We found that when the ACC^D1R^ neurons were irreversibly silenced, the mice became hypersensitive to mechanical stimuli compared to control GFP-expressing mice, as tested using Von Frey filaments and the dynamic brush assay (Fig. 7C, D). When we inactivated the CaMKII ACC neurons using the same strategy, by expressing Kir2.1 to silence and GFP as a control, we found that, in contrast to chemogenetic activation, it caused analgesia to mechanical stimuli in both the baseline and SNI conditions (Fig. S5A-C). Whereas inactivation of the Vgat+ve interneurons with Kir2.1, achieved by expressing Kir2.1 or GFP (controls), in Vgat-Cre mice, resulted in hyperalgesia in baseline and SNI conditions (Fig. S5D-F). In summary, inactivation of the ACC^D1R^ neurons resulted in a similar mechanical hyperalgesia phenotype as in inhibiting the ACC interneurons.

**Figure 7.**
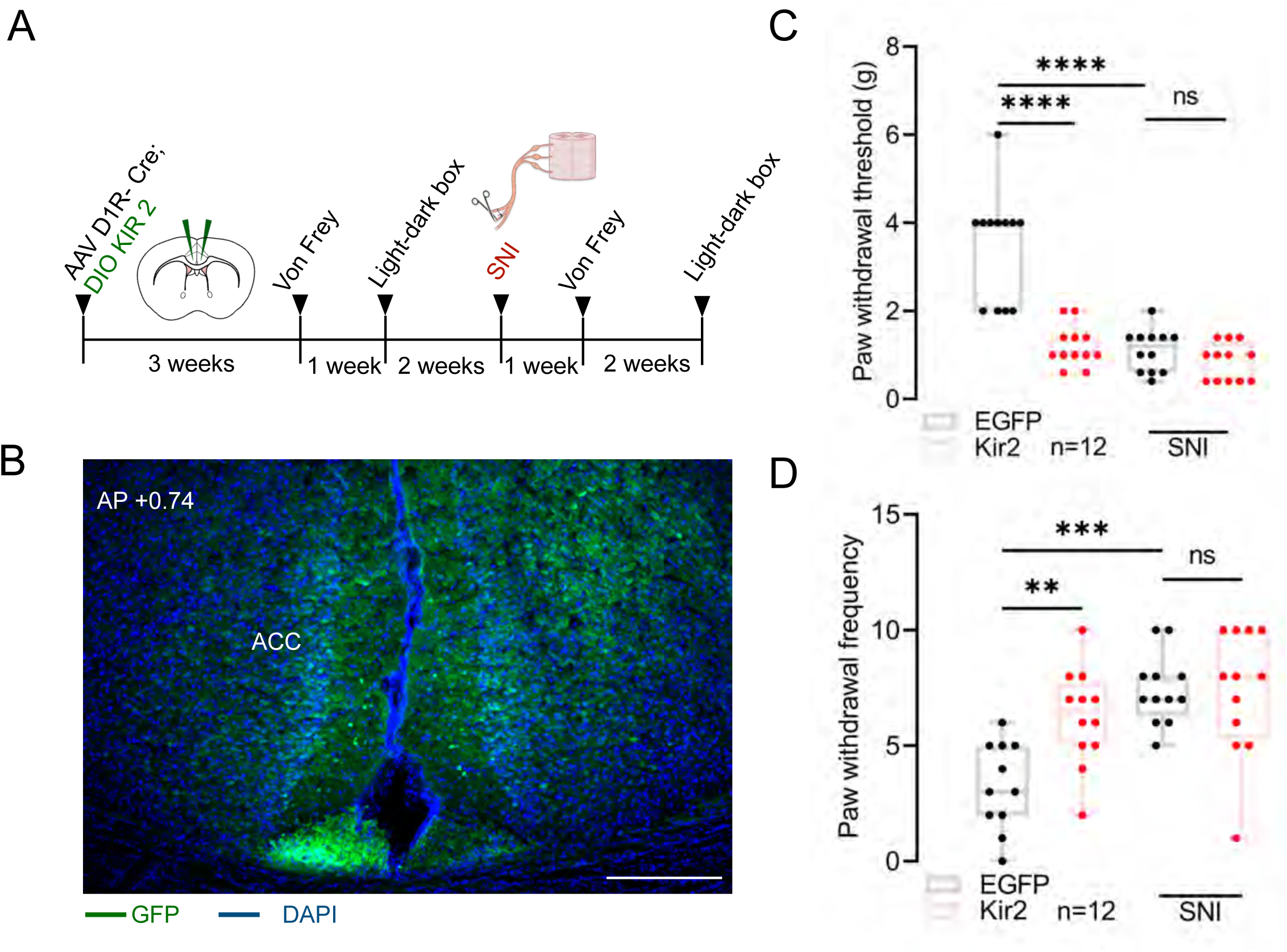
Silencing ACC^D1R^ neurons is sufficient to induce mechanical allodynia. A) Schematic of experimental design. AAV encoding D1R^Cre^ and Cre-dependent kir2 was injected bilaterally into the ACC of CD1 mice. B) Fluorescence microscope image of AAV DIO Kir 2 bilateral expression in ACC D1R neurons. C) Paw withdrawal threshold following inactivation, pre (Tukey’s multiple comparison, 2.300± 0.3006, ****p<0.0001) and post (Tukey’s multiple comparison, p=0.8391) SNI (n=13). D) Paw withdrawal frequency following inactivation pre (Tukey’s multiple comparison, −2.977± 0.8821, **p=0.0082) and post (Tukey’s multiple comparison, p=0.9997) SNI (n=13).

### ACC^D1R^ neurons bidirectionally regulate neuropathic pain-induced negative emotions

In addition to determining mechanical and thermal noxious somatosensory thresholds, ACC has been shown to influence the emotionally aversive components of neuropathic pain[23,31]. Here, we tested whether chemogenetic activation or Kir2.1-mediated silencing of the ACC^D1R^ neurons would influence the anxiety-like behaviors in CD-1 mice with neuropathic pain. We found that expression of hM3Dq-mCherry in the ACC^D1R^ neurons and i.p. administration of DCZ increased the time spent on the lit side of the light-dark box test, indicating a reduction in anxiety (Fig. 8A). The anxiolytic effects of ACC^D1R^ activation persisted in the mice with neuropathic pain, even though mice with SNI displayed heightened anxiety-like behaviors compared to the baseline (Fig. 8A). In contrast, when we silenced the ACC^D1R^ neurons by expressing Kir2.1 under the D1R promoter and used mice with GFP in the ACC^D1R^ neurons as controls, we found that anxiety-like behaviors were increased in the baseline conditions, as well as after SNI (Fig. 8B). Thus, when the activity of the ACC^D1R^ neurons is altered, negative affective-motivational behaviors are promoted, and these emotional effects are exacerbated in animals with neuropathic pain. Since we previously demonstrated that ACC^D1R^ neurons comprise both excitatory and inhibitory neurons (Fig. 3), we next sought to understand which of these two cell populations contributes to the effects of emotional states by ACC^D1R^ modulation. Chemogenetic activation of the CaMKII ACC neurons promoted anxiety-like behaviors by reducing the time spent in the lit side of the light-dark box (Fig. 8C), and Kir2.1-mediated inhibition of the same neurons increased the time spent in the lit side (Fig. 8D). Thus, transient stimulation of ACC pyramidal neurons was anxiogenic, whereas silencing was anxiolytic. Moreover, inactivation of Vgat+ ACC neurons with Kir2.1 channels resulted in enhanced anxiety as observed in mice where ACC^D1R^ neurons were silenced (Fig. S6A, B). Furthermore, we investigated whether chemogenetic activation of the ACC^D1R^ neurons is sufficient to induce a learned preference in the conditioned place preference (CPP) test (Fig. S7A). We found that pairing a chosen chamber of the CPP with stimulation of the ACC^D1R^ neurons by administering DCZ, results in preference for the paired chamber, both in baseline and SNI conditions (Fig. S7B). In contrast, the opposite effect was observed when CaMKII neurons were chemogenetically activated; mice avoided the chamber paired with DCZ (Fig. S7C). Together, our data on the effects of ACC^D1R^ modulation on affective-motivational aspects of neuropathic pain indicate that when both the pyramidal and interneuron sub-population are simultaneously modulated, the effect of modulating the inhibitory neurons may mask the roles of pyramidal cells on affective-motivational behaviors.

**Figure 8.**
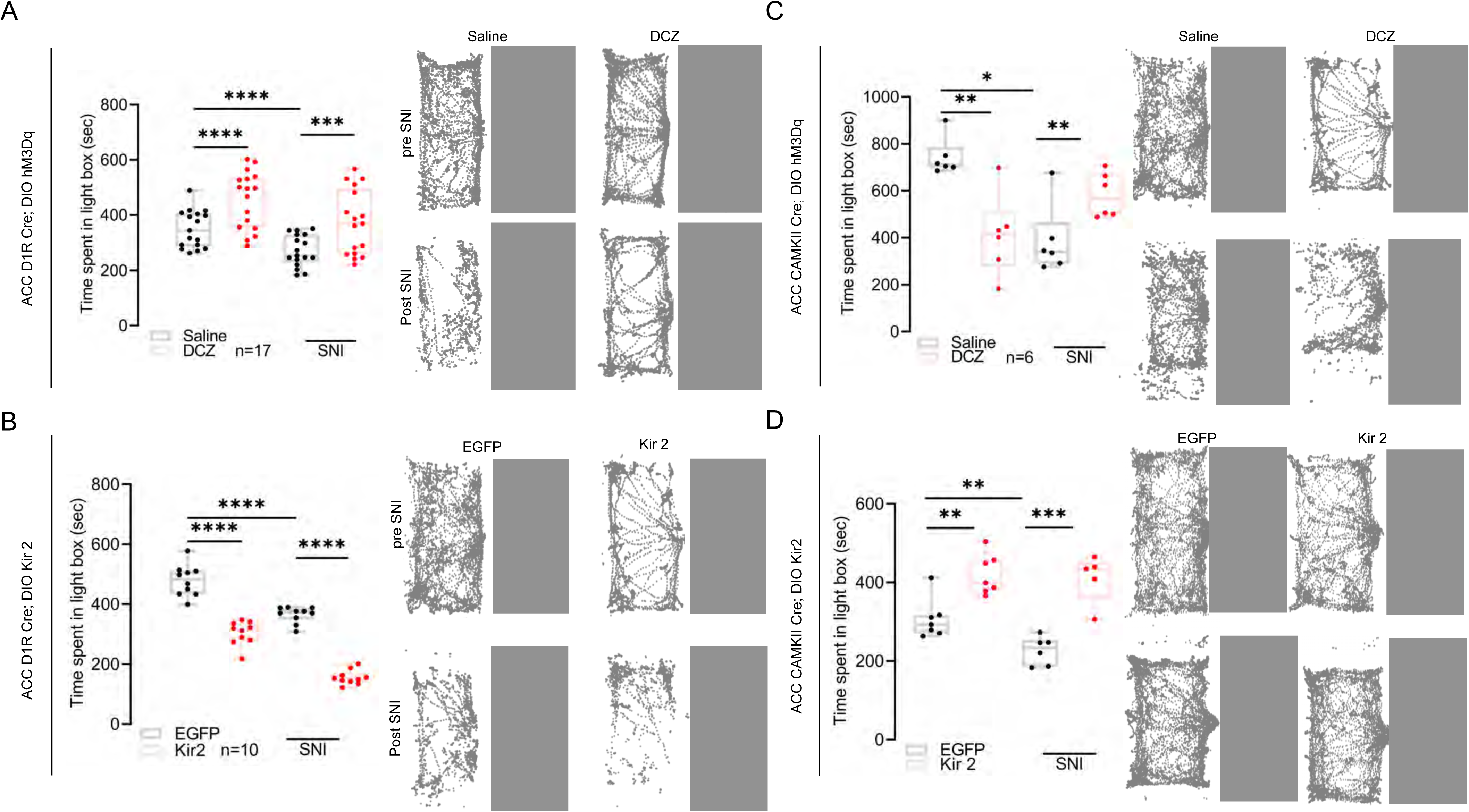
ACC^D1R^ neurons bidirectionally regulate neuropathic pain-induced negative emotions. A) AAV encoding D1R^Cre^ and Cre-dependent excitatory DREADD hM3Dq was injected bilaterally into the ACC of CD1 mice. Time spent in the light box after DCZ administration before (Paired t-test, 1.051±19.47, ****p<0.0001) and after (Paired t-test, 1.02.1± 23.98, ***p=0.0006) SNI (n= 17). On the right side are the trajectory traces of mice navigating through the light-dark arena during saline and DCZ administration before and after SNI. B) AAV encoding D1R^Cre^ and Cre-dependent Kir2 were injected bilaterally into the ACC of CD1 mice. Comparison of time spent in the light box between control and Kir2 mice before (Tukey’s multiple comparison, −174.9± 16.67, ****p<0.0001) and after (Tukey’s multiple comparison, 209.7± 16.67, ****p<0.0001) SNI (n= 13). On the right side are the trajectory traces of control and kir2 mice navigating through the light-dark arena before and after SNI. C) AAV encoding CAMKII Cre and Cre-dependent excitatory DREADD hM3Dq was injected bilaterally into the ACC of CD1 mice. Time spent in the light box after DCZ administration before (Paired t-test, −331.2± 70.14, **p=0.0052) and after (Paired t-test, −194.8± 44.43, **p=0.0071) SNI (n= 6). On the right side are the trajectory traces of mice navigating through the light-dark arena during saline and DCZ administration before and after SNI. D) AAV encoding CAMKII Cre and Cre-dependent Kir2 was injected bilaterally into the ACC of CD1 mice. Time spent in the light box after DCZ administration before (Unpaired t-test, 113.5± 27.09, **p=0.0013, n= 7) and after (Unpaired t-test, 184.5± 29.81, ***p=0.0002, n= 5) SNI. On the right side are the trajectory traces of mice navigating through the light-dark arena before and after SNI.

### Mapping the postsynaptic targets of ACC^D1R^ neurons

Our physiological and behavioral experiments show that the ACC^D1R^ neurons are involved in driving the sensory and affective-motivational pathophysiology associated with neuropathic pain. To understand how ACC^D1R^ neurons engage their downstream targets, we mapped the axonal projections of ACC^D1R^ neurons. To determine the post-synaptic targets of ACC^D1R^ neurons, AAVs encoding D1R-Cre and Cre-dependent tdTomato was injected into the ACC of wild-type CD-1 mice (Fig. 9A). Upon whole brain screening, fore-brain areas such as Medial septum (MS), Claustrum (CLA), Anterior insular cortex (AIC), Caudate putamen (CPu), Basolateral amygdala (BLA), Parafascicular thalamus (PF), Lateral hypothalamus (LHA), and mid-brain areas such as Ventral tegmental area (VTA), Substantia nigra reticulata (SNR), and brainstem nuclei dorsolateral peri aqueductal grey (dlPAG), and Locus coeruleus (LC) were found to be the densely innervated by the ACC^D1R^ neurons. This implies that ACC^D1R^ pyramidal neurons provide inputs to nuclei across the brain while modulating sensory and affective-motivational components of pain.

**Figure 9.**
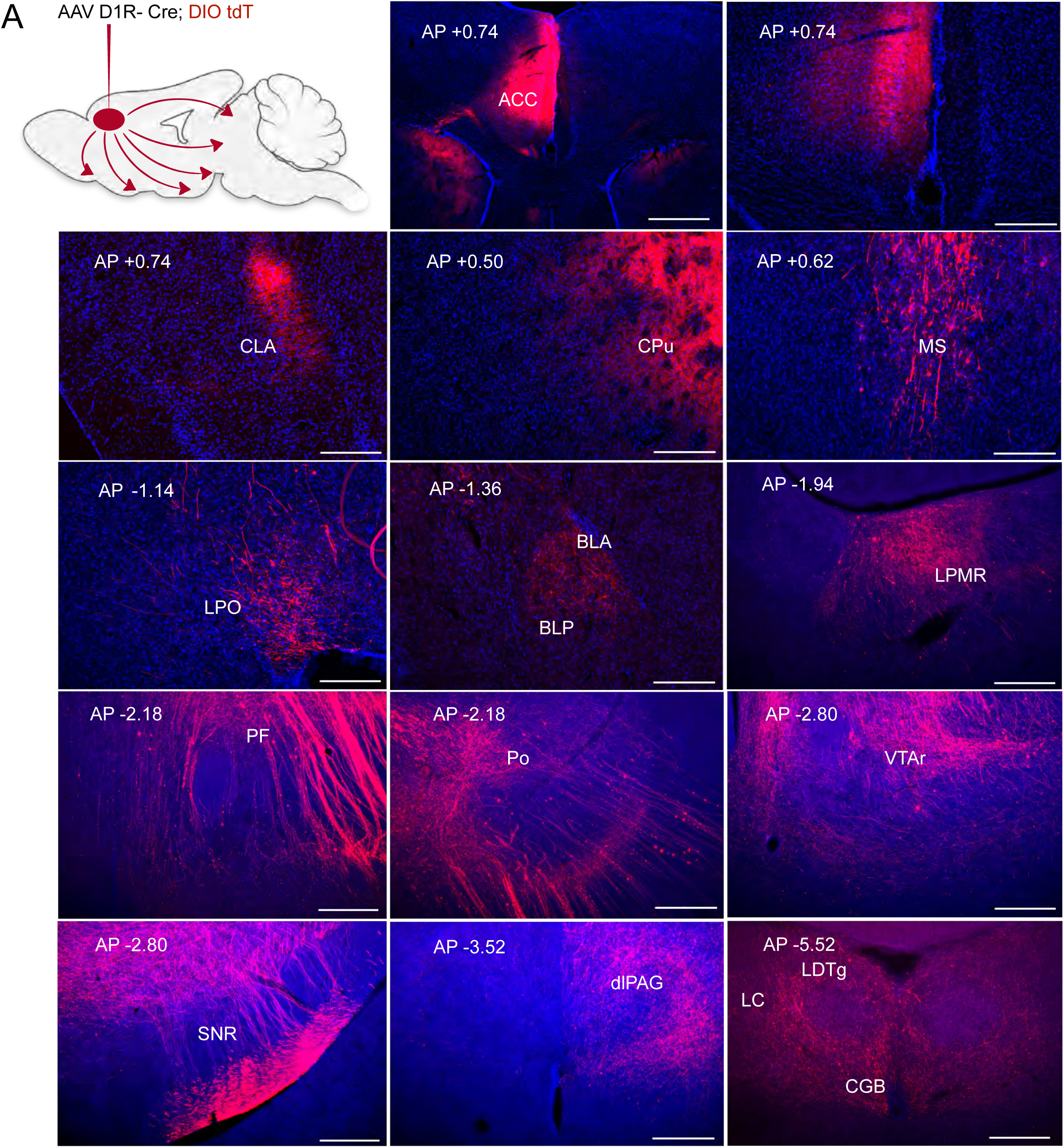
Mapping of ACC^D1R^ inputs with monosynaptic rabies. A) AAV encoding D1R^Cre^ and Cre-dependent TVA and AAV G were injected into the ACC of CD1 mice, and after 21 days, AAV encoding ENVA RV (dG) was injected, and after a week. B) Rabies tracing strategy to study presynaptic partners. C) Fluorescence images showing Rabies labelling in presynaptic regions, Medial septum, Anterior insular cortex, Claustrum, Basolateral amygdala, Parafascicular thalamus, Posterior thalamic nucleus, Lateral hypothalamus, Ventral tegmental area, Substantia nigra, dorsolateral periaqueductal grey. D) Whole brain quantification of the starter cells in ACC along the bregma from +0.10 to +0.26 mm. Data are presented as mean ±SEM, from n=2 mice. E) Whole-brain quantification of the inputs to the ACC^D1R^ neurons. Data are presented as mean ±SEM, from n=2 mice.

### Identification of the brain nuclei projecting to ACC^D1R^ neurons

Thalamocortical pathways are known to be the primary inputs to the ACC through which somatosensory information is relayed[71]. However, most of the neuromodulatory inputs to the cortex arise from distinct midbrain and brainstem structures. Thus, to achieve a comprehensive understanding of how the ACC neurons integrate ascending somatosensory and neuromodulatory inputs, it is imperative that we perform a brain-wide screening for the monosynaptic inputs. To that end, monosynaptic rabies tracing in a Cre-dependent manner can facilitate the mapping of synaptic inputs to molecularly defined neuronal populations in the brain[30,67,68]. We first injected AAVs encoding D1R-Cre and Cre-dependent nls-GFP tagged rabies glycoprotein (RG-nls-GFP) together with TVA (the cognate receptor) in the ACC of CD-1 mice (Fig. 10A, B). After 3 weeks, we delivered EnvA-pseudotyped rabies virus carrying sequences for tdTomato fluorescent protein at the same site (Fig. 10A, B). Upon sectioning and imaging the fluorescence of the whole brain, we found tdTomato in neurons in the forebrain: Medial septum (MS), Claustrum (CLA), Anterior insular cortex(AIC), Basolateral amygdala (BLA), Parafascicular thalamus (PF), Lateral hypothalamus (LHA), mid brain: Ventral tegmental area (VTA), Substantia nigra reticulata (SNR), Substantia nigra per compacta (SNpc) and brainstem dorsolateral peri aqueductal grey (dlPAG) (Fig. 10C). Thus, ACC^D1R^ neurons are reciprocally connected to brain areas known to process nociceptive and somatosensory information, negative affective-motivational states, aversion, and cognition. Interestingly, we found that the VTA and SNR, the primary sources of dopamine in the midbrain, project to the ACC. Hence, we sought to understand the axonal projections of these two nuclei in the ACC. We expressed GFP in the VTA and tdTomato in the SNR/SNpc under the thyroxine hydroxylase (Th) promoter in a Cre-dependent manner using AAVs (Fig. S8A, B). We found that terminals from VTA preferentially terminate in the superficial of the ACC, whereas the SNR/SNpc projections terminate in the deeper layers (Fig. S8B). Since D1R expression is enriched in the deeper layers of the ACC (Fig S3A), the projections of SNR/SNpc to the ACC are of particular interest in modulating pain thresholds and affective-motivational aspects of chronic neuropathic pain.

**Figure 10.**
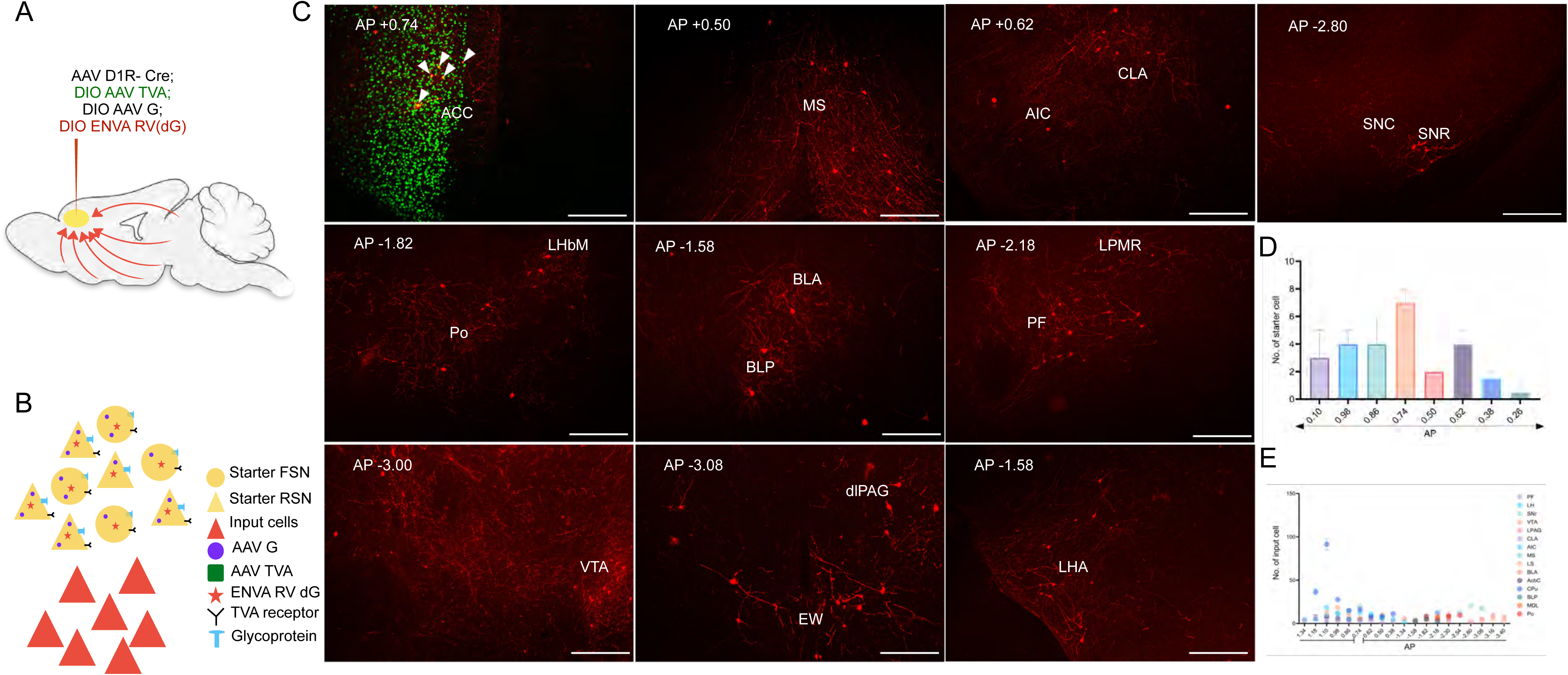
Axonal targets of ACC^D1R^ neurons. A) Schematics showing viral infection with AAVs encoding D1R-Cre and Cre-dependent DIO tdT in ACC in CD1 mice. B) Fluorescence microscope images showing downstream targets, Claustrum, Caudate putamen, Medial septum, Lateral preoptic area, Basolateral amygdala, Parafascicular thalamus, Posterior thalamic nucleus, Ventral tegmental area, Substantia nigra, dorsolateral periaqueductal grey.

## Discussion

Here, we have attempted to address the following questions: a) how dopamine may modulate neuronal activity in ACC to alleviate chronic neuropathic pain, and b) what roles do dopaminergic receptor-expressing neurons in the ACC play in determining the nociceptive thresholds and affective-motivational components of neuropathic pain? From our experiments, we conclude that modulation of D1R receptors can alter the firing properties of ACC pyramidal and interneurons. This modulation was sufficient to reduce mechanical allodynia and anxiety in an animal model of neuropathic pain. When we activated the neurons in the ACC that express D1R, we found that mechanical hypersensitivity and exacerbated anxiety in mice that had undergone SNI were reversed. Whereas inactivation of the same neurons was sufficient to promote mechanical hypersensitivity and anxiety-like behaviors. Our gene-expression studies and electrophysiological data suggest that, although D1R neurons are expressed in both excitatory and inhibitory neurons, when their activity is manipulated simultaneously, the effect of the inhibitory neurons might dominate, in the context of neuropathic pain.

The primary neuromodulatory systems, such as dopaminergic, serotonergic, and noradrenergic inputs to the ACC, have been shown to alter ACC activity and shape behavioral outcomes[34]. Recent findings, as well as our data, suggest that these neuromodulatory systems may change the excitability of ACC neurons, thereby exacerbating or alleviating allodynia and negative affective-motivational states in animal models of neuropathic pain. Here, we propose a mechanism in which D1R receptors are expressed on both pyramidal neurons and interneurons in the ACC, and a reduction in dopamine release due to neuropathic pain has opposing effects on these two types of neurons in the ACC. Simultaneous transient activation of D1R pyramidal and interneurons resulted in increased mechanical thresholds, reduced allodynia, and decreased anxiety. These findings are in agreement with previous observations, where D1R was genetically deleted from all ACC neuronal cell types, resulting in a lowering of mechanical thresholds[16]. We found that this is due to the differential D1R signalling in the pyramidal and interneurons thus influencing the neuronal inhibition and excitation respectively, We found that this is due to the fact that the effect of D1R modulation on pyramidal neurons and interneurons is contrasting, as the application of D1R agonists reduces the excitability of pyramidal neurons and potentiates interneurons (Fig. 2). Even though, the presence of D1R in the ACC interneurons was previously unknown, multiple studies have shown that D1R is enriched in the PV interneurons in the prefrontal cortex of macaques[39], found in the mouse prelimbic prefrontal cortex PV+, VIP+, and SOM+ interneurons[2]and furthermore, a greater proportion of prefrontal cortex GABAergic neurons were found to express D1R than pyramidal neurons[54]

We mapped the inputs and outputs of ACC^D1R^ neurons and found, surprisingly, that these cells are connected bidirectionally to various forebrain, midbrain, and brainstem nuclei, which are known to play important roles in chronic neuropathic pain. For example, multiple studies have reported that the ACC is bi-directionally connected with intercortical nuclei, such as the AIC and Claustrum[41,63,70], as well as subcortical structures, including the LHA and BLA[4,36]. Through lesion studies in human subjects, AIC has been hypothesized to tune cortical areas to cognitive information during afferent processing[60]. Furthermore, AIC neurons were found to be sufficient to induce hyperalgesia and aversion in mice[11]. Thus, mechanistic studies dissecting the role of reciprocal circuitries between AIC and ACC can reveal unique insights into how emotional and cognitive information shape pain outcomes. A recent study found direct orexigenic LHA inputs to the ACC driving stress-induced anxiety[36]. Given the roles played by LHA in the interactions between stress and pain[58], it would be interesting to test whether the affective-motivational behaviors mediated by ACC^D1R^ neurons are affected by LHA inputs. Interestingly, we found that the SNR/SNpc project to the deeper layers of the ACC (Fig. S8A, B), where the density of the D1R is the highest, implying that D1R-mediated dopaminergic modulation in the ACC primarily occurs through SNR/SNpc inputs, rather than the VTA. Comparative expression of D1R and D2R in the ACC may reveal if the neurons that express these receptors are differentially or preferentially targeted by the midbrain dopaminergic nuclei, VTA and SNR/SNpc. The fact that chronic neuropathic pain is often a non-motor symptom in patients with Parkinson’s disease[18], SNR/SNpc projections to the D1R-rich layer in the ACC can have significant implications.

There are a few technical and conceptual drawbacks of this study that need to be addressed in the future to unravel the role of dopamine signaling in allodynia and anxiety observed due to neuropathic pain. Recent advancements in intersectional optogenetics[22,51], combined with transgenic mouse strains expressing Cre under dopamine receptor promoters, such as D1R and D2R, can help further elucidate how dopamine receptors and receptor-expressing neurons in the ACC mediate the pathophysiology of neuropathic pain. AAV-based CRISPR-CAS9[26] systems can help us further understand how knocking out the dopamine receptors, and thus the downstream signaling may regulate allodynia, hyperalgesia, and pain-induced anxiety. Knocking out the receptors genetically may provide us with unique insights into how dopaminergic modulation regulates ACC function. Here, we have focused on the SNI model of neuropathic pain in mice; however, it will be important to study whether the dopaminergic modulation of ACC neurons underlies allodynia, hyperalgesia, and pain-induced anxiety observed in other neuropathic models, such as chemotherapy drug-induced neuropathy, diabetic neuropathy, or peripheral inflammatory pain[32]. Furthermore, genetically encoded dopamine sensors[13,46,53], in combination with calcium indicators, will help us gather real-time information on how dopamine and neural dynamics correspond with animal behavior in neuropathic pain.

## Materials and Methods

### Animals

Animal care and experimental procedures were performed per the protocol approved by the CPSCEA at the Indian Institute of Science. The ethics approval number is CAF/ETHICS/988/2023. The animals were housed at the IISc Central Animal Facility under standard animal housing conditions: a 12-hour light/dark cycle from 7:00 am to 7:00 pm, with ad libitum access to food and water. Mice were housed in IVC cages in Specific Pathogen-Free (SPF) clean air rooms. Mice strains used: CD1 mice (Jackson Laboratories, USA). Vgat-ires-cre knock-in (C57BL/6J) or Slc32a1-IRES-Cre[65] was purchased from Jackson Laboratortories, USA and bred with CD-1 to obtain uniform genetic background with the control animals. The experimental animals were between 6 and 8 weeks old.

### Stereotaxic injections

Mice were anaesthetised with 2% isoflurane/oxygen before and during the surgery. A craniotomy was performed at the marked point using a handheld microdrill (RWD, China). A Hamilton syringe (10 μL) with a glass-pulled needle infused 150 nL of viral particles (1:1 in saline) at 100 nL/minute. The following coordinates were used to introduce virus/dyes: ACC -Anterior-Posterior (AP): +0.74 mm, Medial-Lateral (ML): + 0.25 mm; Dorsal-Ventral (DV): −1.50 mm; VTA -AP: −3.08 mm, ML: ±0.50 mm; DV: −4.35 mm. After each injection, the needle was left in place for a few minutes to allow the virus to diffuse before withdrawal. The scalp was then sutured, and mice were monitored for 30 minutes post-surgery. Meloxicam (I.P.) (10mg/25g) was administered to minimise postoperative pain and inflammation.

### Viral vectors

Chemogenetic activation and inhibition: rAAV-D1(SP)-CRE-WPRE-bGH pA (Brain VTA, Catalog# PT-0570), pAAV-hsyn-DIO-hM3D(Gq)-mCherry (addgene, Catalog # 44361, titer-1.8 x 1013 GC/ml), rAAV-EF1α-DIO-Kir2.1-P2A-EGFP WPRE-hGH polyA (Brain VTA, Catalog# PT- 1401, titer- 2 x 1012 vg/ml)

VTA DAT knockout: rAAV-mTH-NLS-CRE-WPRE-SV40 polyA (Brain VTA, Catalog# PT- 7081), rAAV-CMV-FLEX-NLS-hSaCas9-NLS-3XHA-bGH-polyA-U6-sasgRNA(Slc6a3)(Brain VTA, Catalog# PT- 10092), rAAV-CMV-FLEX-NLS-hSaCas9-NLS-3XHA-bGH-polyA-U6-sasgRNA(scramble) (Brain VTA, Catalog# PT- 6643)

Neuronal activity: rAAV-D1(SP)-CRE-WPRE-bGH pA (Brain VTA, Catalog# PT- 0570), AAV9-Syn-FLEX-jGCaMP8s-WPRE (Addgene, Catalog# 162377)

For dopamine sensing in ACC and VTA: rAAV-D1(SP)-CRE-WPRE-bGH pA (Brain VTA, Catalog# PT- 0570), rAAV-mTH-NLS-CRE-WPRE-SV40 polyA (Brain VTA, Catalog# PT- 7081), rAAV-EF1α-DIO-dLight1.2-WPRE-pA (Brain VTA, Catalog# PT- 1296)

Chemogenetic controls: AAV_ pCAG-FLEX-Egfp-WPRE (Addgene, Catalog# 51502), pAAV-FLEX-tdTomato (Addgene, Catalog# 28306, titer- 1.6 x 1013 GC/ml)

Mapping VTA and SNr dopaminergic projections: rAAV-mTH-NLS-CRE-WPRE-SV40 polyA (Brain VTA, Catalogue # PT-7081), AAV-pCAG-FLEX-Egfp-WPRE (Addgene, Catalogue # 51502), pAAV-FLEX-tdTomato (Addgene, Catalogue # 28306). Post-hoc histological examination of each injected mouse was used to confirm that viral-mediated expression was restricted to target nuclei.

### Spared nerve injury surgery

Spared nerve injury (SNI) surgery was performed as previously described[69], inaccordance with the guidelines of the International Association for the Study of Pain (IASP) on the use of awake animals in pain research. The mice (6 weeks old) were anaesthetised with ketamine (100 mg/kg) and xylazine (15 mg/kg) administered intraperitoneally (0.1 mL per 20 g body weight). Body temperature was maintained at 37 °C using a heating pad (HBLD-01, Orchid Scientific). Following shaving of the left lateral thigh, a skin incision was made, and the femoral muscle was carefully dissected, exposing the sciatic nerve and its three terminal branches: the common peroneal (CPN), tibial (TN), and sural nerves. The CPN and the TN were ligated using an absorbable polyglactin surgical suture and transected distally to prevent regeneration, thereby leaving the sural nerve intact. The skin was sutured with a nylon surgical suture. Meloxicam was administered postoperatively to minimise pain and prevent infection.

### Preparation of ACC slices

Mice were deeply anesthetized with isoflurane and transcardially perfused with ice-cold, oxygenated (95% O2/5% CO2) sucrose-containing artificial cerebrospinal fluid (s-aCSF) containing (in mM): 75 NaCl, 25 NaHCO3, 15 dextrose, 75 sucrose, 1.25 NaH2PO4, 2 KCl, 2.4 Na pyruvate, 1.3 L-ascorbic acid, 0.5 CaCl2, 3 MgCl2, with an osmolarity of 290-295 mOsm/L. Following perfusion, the brain was quickly removed, and 300-µm-thick coronal slices of the anterior cingulate cortex were prepared in ice-cold s-aCSF using a vibratome (VT1200S, Leica). The slices were then incubated at 33°C for 30 minutes in oxygenated ACSF containing (in mM): 126 NaCl, 26 NaHCO3, 10 glucose, 1.25 NaH2PO4, 3 KCl, 2 CaCl2, and 1 MgCl2, with an osmolarity of 305-310 mOsm/L. After incubation, slices were allowed to recover in oxygenated ACSF at room temperature for 1 hour before recording.

### Whole cell electrophysiology

Slices were transferred to the recording chamber of an Olympus BX51W1 upright microscope and continuously perfused with oxygenated artificial cerebrospinal fluid (aCSF) via a peristaltic pump (2 mL/min). EGFP-labelled ACC D1R neurons were visualised using a CoolLED pF-300 lite epifluorescence source. To assess the firing properties of ACC neurons, current clamp recordings were obtained from visually identified ACC neurons using patch electrodes (4-8 MV) filled with intracellular solution containing (in mM): 145 K-gluconate, 4 NaCl, 10 HEPES, 14 Na-phosphocreatine, 4 Mg ATP, 0.3 Na GTP, and 3 L-ascorbic acid with an osmolarity of 285-295 mOsm/L. In the current-clamp configuration, incremental current steps of 20 pA (from −20 pA to 260 pA) were applied. Following current clamp recordings, spontaneous excitatory postsynaptic currents (EPSCs) were recorded in a voltage clamp configuration with cells held at −70 mV.

For whole cell pharmacology recordings with SKF 83822, we used 10uM concentration after 20 mins of bath application to acquire action potential recordings. Control versus drug patch clamp experiments were done intermittently, sometimes on the same day.

For whole cell paired LTP induction stimulation electrode was kept in the layer ⅚ near the cell to be patched and recordings were performed in voltage-clamp mode and cells were voltage-clamped at ∼-70mV mV. The stimulation intensity was chosen such that it evoked 10-20% of the maximum excitatory postsynaptic currents (EPSCs) response. LTP was induced using a pairing protocol where a train of pulse stimulation at 1.4 Hz was paired with a depolarizing step to 0 mV for 125 seconds[28]. EPSCs were recorded for 20-30 min after the LTP induction.

Data were acquired using a Multiclamp 700B amplifier and pClamp 10.5 software (Molecular Devices), filtered at 2 kHz, and digitised at 20 kHz using a Digidata 1550 data acquisition system (Molecular Devices). The electrophysiology data were analysed using Clampfit.

### For ACC field-electrophysiology recordings

Slices were transferred to the recording chamber on an Olympus SZ51 stereomicroscope, and oxygenated aCSF was being perfused by a peristaltic pump (Watson-Marlow IP31) at a rate of 4.5 ml/min. For the acute slice stimulation experiments, an FHC cluster electrode was placed in layer 1-2 of the ACC, and the recording electrode was placed in layer 5-6. The electrical stimulation pulse was generated using an Isoflex stimulator box by AMPI as described previously [1]. For recording the postsynaptic response, an aCSF-filled glass pipette with an A-M Systems microelectrode holder was used. The signal was amplified using A-M Systems’ microelectrode AC amplifier and digitized with the National Instruments data acquisition system (NI PCI-6221).

The magnitude of pulse stimulation intensity was set to 50% of the stimulation causing maximum fEPSP response for each slice. For baseline LTP experiments in ACC slices, single-pulse electrical stimulation was initially applied for 15 minutes to record stable fEPSP responses. LTP was induced using 100 Hz high-frequency stimulation (HFS), repeated four times with an ∼1-second interval between each high-frequency sweep. Post-LTP fEPSP responses were recorded using single pulse stimulation for at least 45 minutes to measure long-term synaptic changes.

For LTP drug experiments, baseline fEPSP responses were recorded for 10 mins in normal aCSF (mixed with vehicle control if needed), followed by 10 mins of recordings in aCSF mixed with 10 μM SCH (SCH 23390 hydrochloride, Cat. # 0925, Tocris Bioscience) or 50 μM SKF (Tocris Bioscience). LTP was induced using 100 Hz HFS repeated four times as stated above. Drug washout was performed for 2 mins with normal aCSF, and post-LTP responses were recorded for at least 45 mins in the absence of drug. The recording and analysis workflow was done in WinLTP software (version 1.10) from the University of Bristol.

### Intracranial ACC cannulation

Stereotaxic cannulation surgeries were performed in mice under continuous anesthesia with 2% isoflurane/oxygen before and during the surgery. Unilateral cannulation of the ACC was performed. A pair of 26-gauge stainless steel guide cannulas, cut 6 mm below the pedestal, was implanted targeting the ACC injection site: AP, +0.74; ML, ±0.25 mm; DV, −1.50 mm. Guide cannula was affixed to the skull using dental cement. A stainless-steel dummy cannula was inserted into each guide to keep the cannula free of debris. Mice were allowed to recover for 2 to 3 days. Cannula placement was verified post hoc by histological examination.

### SKF 83822 dosage and administration

For behavioral assays, the D1R agonist SKF 83822 (Cat. # 2075, Tocris Biosciences) was dissolved in saline to yield a working concentration of 4 µg/2 µl. A volume of 200 nl was infused into the anterior cingulate cortex (ACC) at a rate of 100 nl/min using the RWD infusion system. Following infusion, the needle was left in place for 5 minutes to allow for drug diffusion, after which the animals were returned to their cages. Behavioral testing commenced 20 minutes post-infusion.

For electrophysiological recordings, SKF83822 was dissolved in DMSO to make a 10 µM working solution. ACC brain slices were incubated in the drug solution throughout the duration of whole-cell patch clamp recordings.

### Behavioural assays

A single experimenter handled behavioural assays for the same cohorts. Before testing, mice were habituated in their home cages for at least 30 minutes within the behaviour testing room and randomized for mouse genotypes and AAV injections. Experimental cohorts maintained an equal male-to-female ratio unless otherwise noted, with no significant sex differences observed in behavioural responses. Throughout the study, every effort was made to minimise animal use.

### Mechanical hyperalgesia and allodynia test

Mice were habituated in a transparent Plexiglass chamber (16 x 16 x 16 cm) placed on an elevated wire mesh platform before testing. Mechanical sensitivity was assessed using a calibrated series of von Frey filaments (0.4 g to 8.0 g), applied perpendicularly to the plantar surface of each hind paw until the filament bent (within 3 seconds). A positive nociceptive response was defined as paw withdrawal, licking, flinching, or shaking. Testing began with a 1 g filament; based on the animal’s response, higher or lower filaments were applied accordingly. The filament strength that results in a 50% likelihood of eliciting a withdrawal response was used as the paw withdrawal threshold. This value serves as an indicator of mechanical allodynia development.

### Dynamic brush assay

Mice were habituated for 15 minutes in a transparent chamber (16 x 16 x 16 cm) placed on an elevated wire mesh platform before testing. Mechanical stimulation was delivered by gently brushing the plantar surface of the hind paw with a soft brush, applying ten strokes along the length of the paw while keeping the brush perpendicular to the paw surface. A positive nociceptive response was defined as paw withdrawal, licking, flinching, or shaking. The relative change in the response was considered an indicator of mechanical allodynia development.

### Light-dark box assay

Anxiety-like behaviour was assessed using a light-dark box apparatus consisting of two chambers (light and dark), measuring 40 cm (length) x 20 cm (width) x 36 cm (height). The chambers were connected by a small opening, allowing mice to access one from another freely. Mice were placed in the light chamber at the start of the 15-minute test and allowed to explore freely. Behaviour was recorded from above using a Logitech C930e camera. Time spent in the light chamber was used as a measure of anxiety-related behaviour. The apparatus was thoroughly cleaned with 0.1% acetic acid between trials. The movement of the mouse was tracked and analysed using DeepLabCut.

### Conditioned place preference and/ or aversion test

A custom-built three-compartment CPP apparatus was used to assess conditioned place preference or aversion (CPP) in mice. The two outer chambers measured 32 cm (length) x 32 cm (width) x 28 cm (height), and the middle neutral chamber measured 11 cm x 8.5 cm. One outer chamber featured white striped walls and a steel mesh floor, while the other had black walls and a steel rod floor. The neutral middle chamber had grey walls and a smooth PVC floor. Entry between chambers was controlled by two manual doors that could be closed to isolate any section. Light intensity was kept constant to avoid innate chamber preference. Chambers were cleaned with 70% ethanol between trials. Mouse activity was recorded using a Logitech C930e camera.

The unbiased CPP/A protocol spanned five days. On Day 1 (pre-conditioning), mice were placed in the central chamber and allowed to freely explore all compartments for 15 minutes; time spent in each outer chamber (T1 and T2) was recorded. Mice with a baseline preference ratio (T1/T2) between 2:3 and 3:2 were considered for conditioning; those outside this range were excluded to ensure unbiased baseline preference.

During Days 2-4 (conditioning phase), mice underwent twice-daily conditioning sessions (morning and evening), separated by at least four hours. On Day 2, the striped chamber was paired with an i.p. injection of DCZ, and mice receiving i.p. saline were confined to the black chamber for 15 minutes. In the evening, mice received i.p. Saline and were confined to the black chamber for 15 minutes. On Day 3, the pairing was reversed (morning saline in black chamber; evening DCZ in striped chamber). On Day 4, the original Day 2 sequence was repeated.

On Day 5 (post-conditioning), both doors of the chamber were opened, and mice were allowed to explore freely for 15 minutes. Time spent in the DCZ-paired chamber was compared between the pre-conditioning (Day 1) and post-conditioning sessions (Day 5) to assess the development of conditioned place preference/aversion.

### DeepLabCut for tracking mice

The tracking of mice in the light-dark box test and the CPA test was done using DeepLabCut (DLC) (version 2.3.9)[35,38]. Data were processed and analysed on a custom-built workstation equipped with an AMD Ryzen 9 5900X 12-core processor and an NVIDIA GPU. For training the DeepLabCut (DLC) model, 30 frames were manually labelled from each of the five videos. Following training, the model was used to analyse the videos, generating position plots and corresponding output in .csv format. For visualisation, representative plots of tracked spine positions were produced.

### Immunohistochemistry and multiplex fluorescent in situ hybridisation

Mice were anaesthetised with isoflurane and perfused transcardially with 1X phosphate-buffered saline (PBS; Takara catalogue # T9181) and 4 % paraformaldehyde (PFA; Ted Pella, Inc. catalogue # 18505). Harvested brains were further fixed in 4% PFA overnight and subsequently transferred to 15% and 30% sucrose for serial dehydration. Brain tissues were placed in the Cryo-Embedding Compound (Ted Pella, Inc.) and frozen at −40°C. Subsequently, 50 µm-thick coronal brain sections were cut using a cryostat (RWD Minux FS800).

For immunostaining experiments, tissue sections were rinsed in 1X PBS (3 times) and incubated in the blocking buffer (5 % Bovine Serum Albumin (BSA) + 0.5 % Triton X-100 + 1X PBS) (BSA-HIMEDIA catalogue # TC194, Triton X-100 SRL catalogue # 64518) for one hour at room temperature. Sections were then incubated in the primary antibody solution at 4℃ overnight. Following three washes in 1X PBS + 0.5% Triton X-100 solution, sections were incubated for 2 hours in an Alexa Fluor-conjugated secondary antibody solution, along with DAPI nuclear stain (SRL catalogue #18668). After washing, sections were mounted onto charged glass slides (Ted Pella, Inc., catalogue # 260382-3) and cover-slipped with Citifluor AF-1 mounting media (Ted Pella, Inc., catalogue # 19470-1) using blue star microscopic cover glass (24 x 60 mm, 10 G). Fluorescence imaging was performed on an upright fluorescence microscope (Khush Enterprises, Bengaluru) using 2X, 4X, and 10X objectives, as well as on a Confocal Microscope (Leica SP8 Falcon, Germany), at the Division of Biological Sciences, Core Imaging Facility, IISc. Image processing was conducted using ImageJ/FIJI processing software, and the confocal images were also analyzed using the Leica Image Analysis Suite software.

### Primary and Secondary antibodies

Primary antibodies: Chicken anti-GFP (1:500 solution; aveslabs catalogue # 1010), Rabbit monoclonal anti-Phospho-c-Fos (Ser32) (1:500 dilution; Cell Signalling Technology Catalogue # 5348), Goat anti-tdTomato (1:1000 dilution; SICGEN catalogue # AB8181).

Secondary antibodies: Goat anti-Chicken IgY (H+L), Alexa Fluor™ 488 (1:1000 dilution; Invitrogen catalogue # A11039), Donkey anti-Rabbit IgG (H+L), Alexa Fluor™ 488 (1:1000 dilution; Invitrogen catalogue # A21206), Donkey anti-Goat IgG (H+L), Alexa Fluor™ 594 (1:1000 dilution; Invitrogen catalogue # A11058).

### Quantification and statistical analysis

Statistical comparisons were performed using the one-way ANOVA test, followed by Tukey’s multiple comparison test and Student’s t-test with GraphPad Prism 8.0.2. For pharmacological activation of D1R receptor behaviour analysis we used two-tailed unpaired t-tests. For analyzing ACC field LTP experiments and pharmacology data unpaired t-test was used. For whole cell electrophysiology data two-tailed unpaired t-test was performed. For chemogenetic activation and inactivation behavioral data Tukey’s multiple comparison test and two-tailed paired test was performed. Data are means, and error bars represent the standard error of the mean (SEM) as indicated. Statistical significance is represented as p>0.05 (ns), p<0.05 (∗), p<0.01 (∗∗), p<0.001 (∗∗∗), p ≤ 0.0005 (∗∗∗∗).

---

**Supplementary Figure 1:**
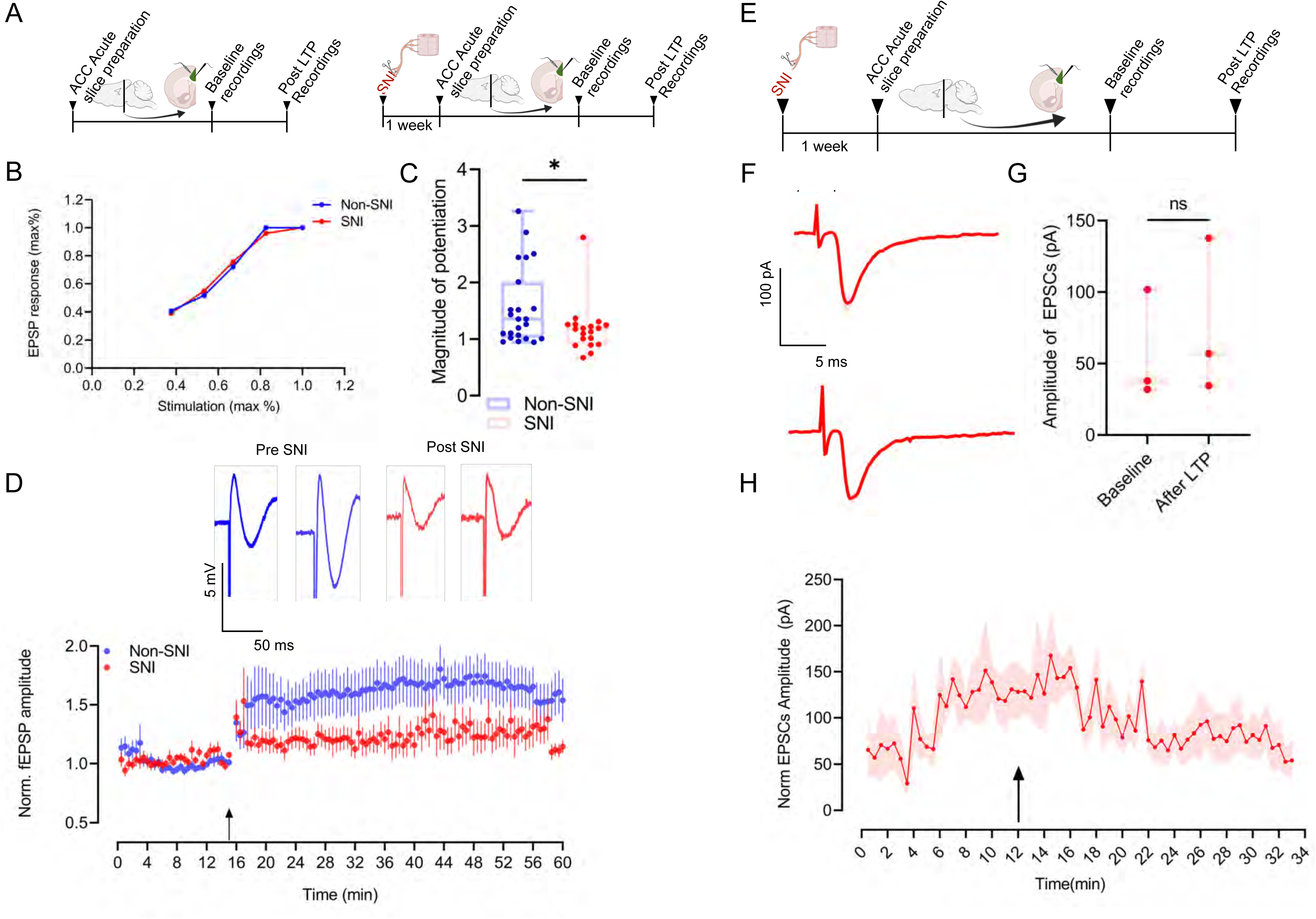
related to- Fig 2. Synaptic plasticity in ACC is impaired following chronic pain and is dopamine-dependent. A) Schematics of field recordings from layer ⅚ in ACC in basal (Non-SNI) and SNI condition. B) Input-output curve. C) Comparison of the magnitude of potentiation after 100 Hz HFS in basal (n=23) and SNI (n=19) conditions. Unpaired t-test, 0.3827±0.1819, *p=0.0417. D) Time course plot showing normalized fEPSPs amplitude with 15-minute baseline and followed by 45 minutes recordings post 100 Hz HFS in basal and SNI condition. Representative traces showing fEPSP amplitude before and after LTP induction in basal and SNI conditions respectively. E) Schematics of paired LTP induction paradigm in whole cell configuration from layer ⅚ in acute ACC slices retrieved from mice transfected with AAV encoding D1R^Cre^ and Cre-dependent eGFP. F) Representative traces showing EPSCs amplitude before and after LTP induction in SNI conditions. G) Comparison of magnitude of potentiation in SNI condition (n=3), Paired t-test, p= 0.1834. H) Normalized EPSC amplitude of ACC^D1R^ neurons in the SNI condition with 12 minutes baseline and followed by 20 minutes recordings post paired LTP induction.

**Supplementary figure 2:**
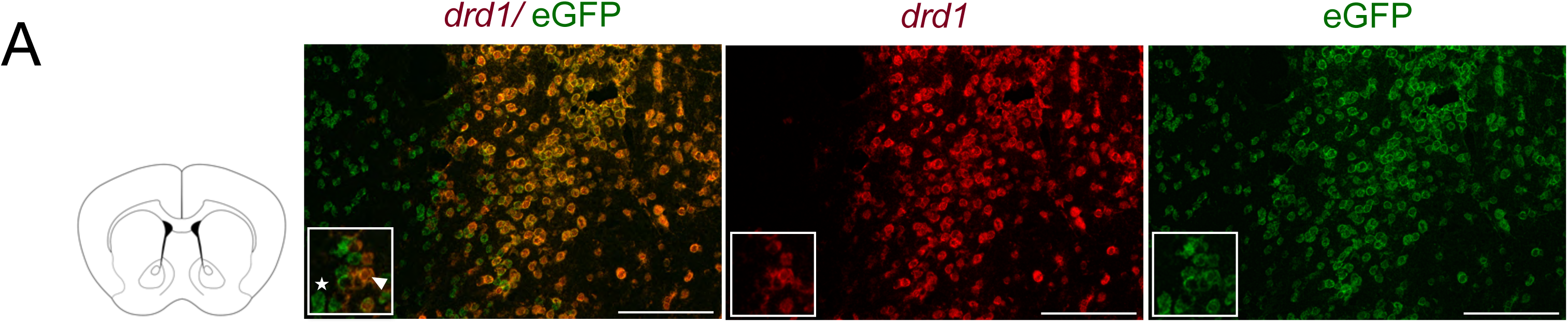
related to- Fig 3. D1R-expressing ACC neurons are a heterogeneous population of excitatory and inhibitory neurons. A) Multiplex in-situ hybridisation, NAc sections were recovered from mice injected with AAV D1R^Cre^ DIO eGFP and probed for drd1 and eGFP, showing the efficacy of the AAV virus.

**Supplementary Figure 3:**
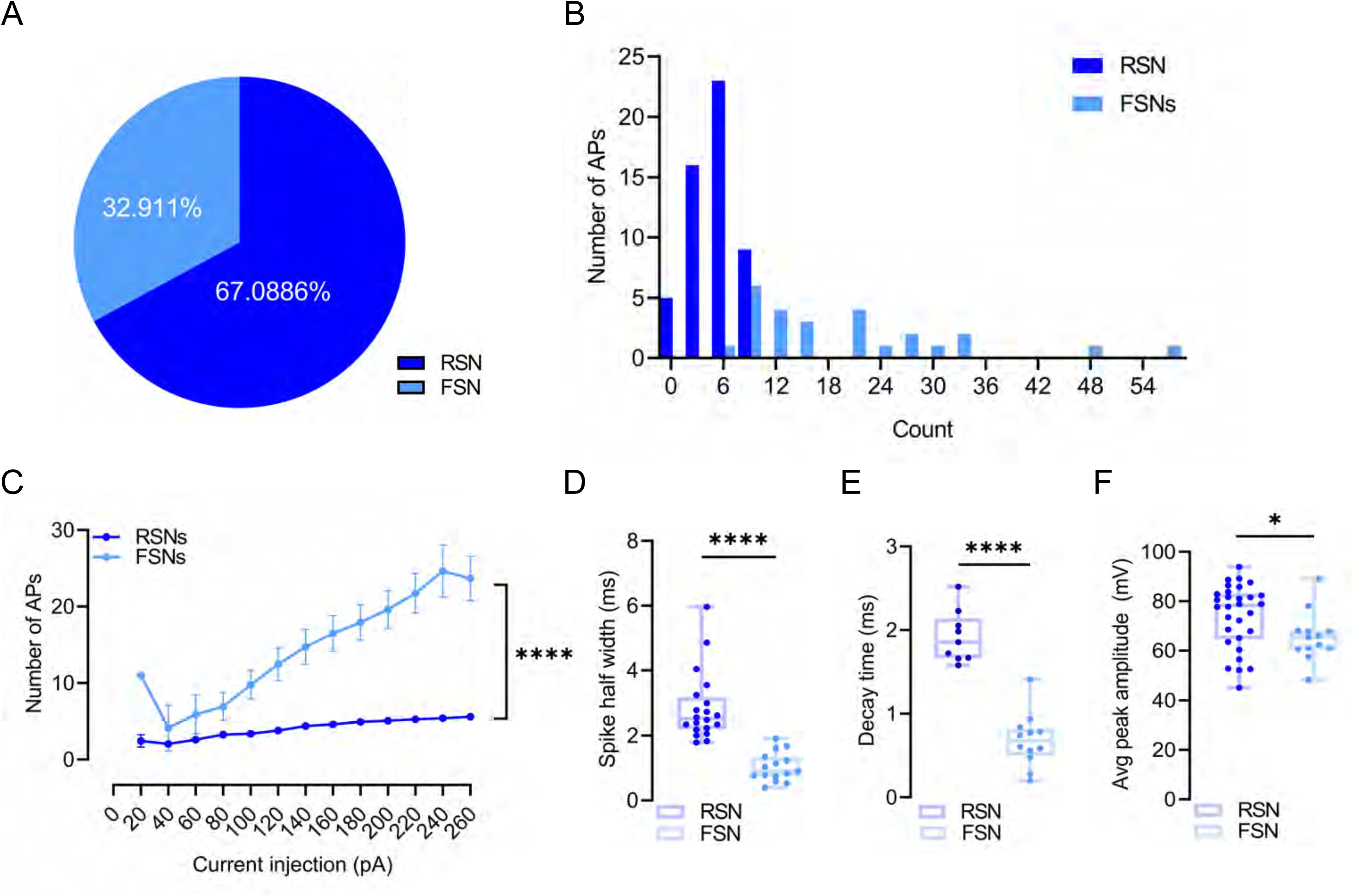
related to- Fig 4. Nerve injury-induced potentiation of ACC neurons can be scaled down with D1R agonist in ex vivo recordings. A) Pie chart showing the RSN and FSN distribution among the ACC neurons patched during whole cell patch clamp electrophysiology recordings. B) Histogram showing frequency distribution of number of action potentials among RSN (n=53), and FSN (n=26). C) Comparison of action potential firing of RSN (n=53), and FSN (n=26) ACC^D1R^ neurons during baseline, Unpaired t-test, 3.680±1.574, *p= 0.0073, 6.380±1.494, ****p< 0.0001, 8.656±1.649, ****p< 0.0001, 10.38±1.728, ****p< 0.0001, 11.89±1.720, ****p< 0.0001, 3.00±1.694, ****p< 0.0001, 14.56±1.765, ****p< 0.0001, 16.48±1.867, ****p< 0.0001, 19.23±2.405, ****p< 0.0001, 18.07±2.072 ****p< 0.0001. C) Comparison of APs spike half-width of RSN (n=20) and FSN (n=15) ACCD1R neurons, Unpaired t-test, 1.821±0.2910, ****p<0.0001. E) Comparison of APs decay time of RSN (n=9) and FSN (n=12) ACC^D1R^ neurons, Unpaired t-test, −1.236±0.1384, ****p<0.0001. F) Comparison of APs spike amplitude of RSN (n=30) and FSN (n=13) ACC^D1R^ neurons, Unpaired t-test, 8.522±4.019, *p= 0.0401.

**Supplementary Figure 4:**
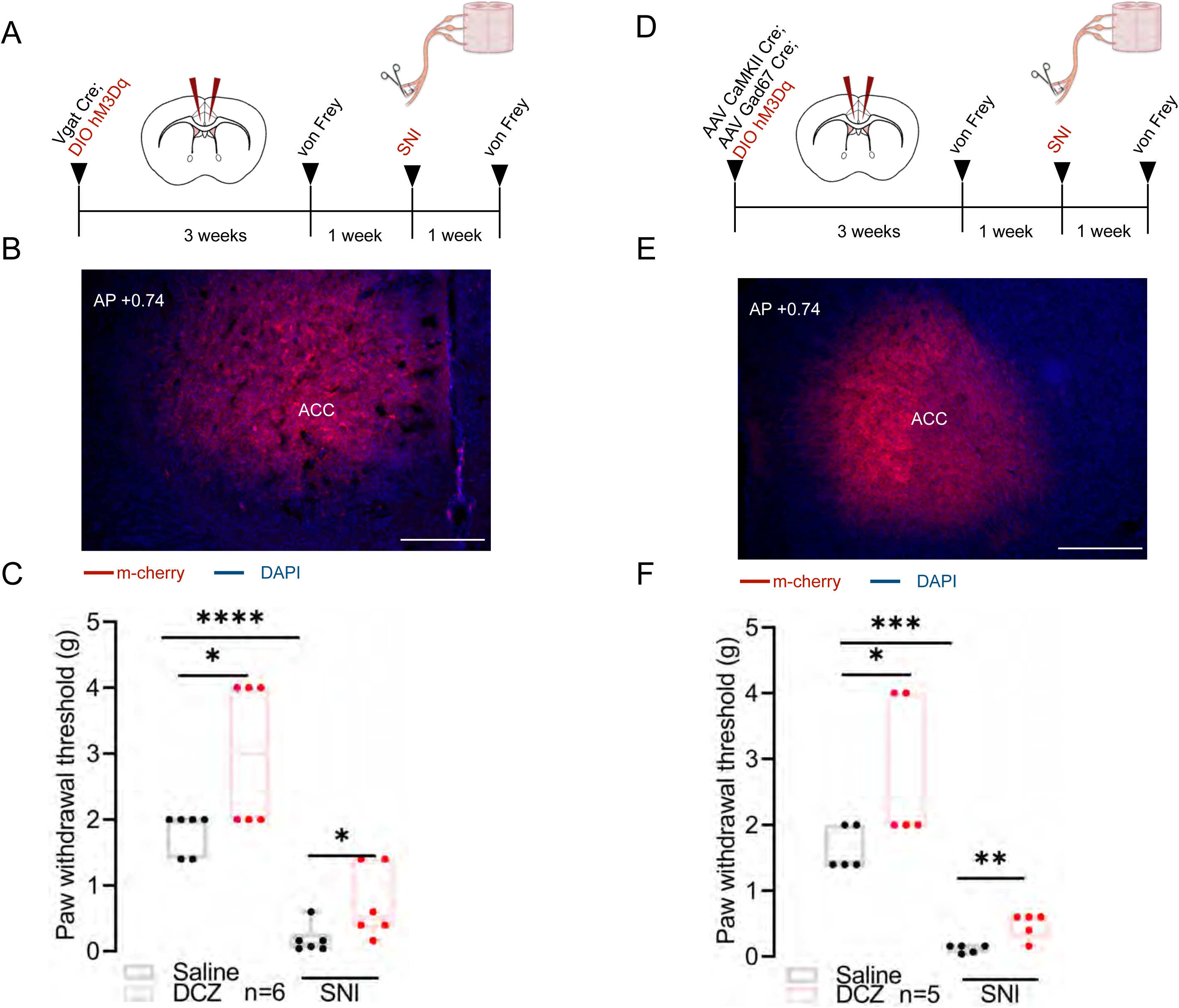
related to- Fig 6. Activating ACC^D1R^ neurons alleviates SNI-induced mechanical allodynia. A) Schematic of experimental design. AAV encoding excitatory DREADD hM3Dq was injected bilaterally into the ACC of Vgat Cre mice. B) Fluorescence microscope image of AAV DIO hM3Dq bilateral expression in ACC Vgat neurons. C) Paw withdrawal threshold pre (Paired t-test, 1.200± 0.3688, *p=0.0226) and post (Paired t-test, 0.5483± 0.1668, *p=0.0218) SNI upon DCZ administration (n=6). C) Schematic of experimental design. AAV encoding CAMKII Cre and Gad 67 Cre and Cre-dependent excitatory DREADD hM3Dq was injected bilaterally into the ACC of CD1 mice. D) Fluorescence microscope image of AAV DIO hM3Dq bilateral expression. E) Paw withdrawal threshold pre (Paired t-test, 1.1160± 0.3429, *p=0.0277) and post (Paired t-test, 0.3540± 0.06226, **p=0.0047) SNI upon DCZ administration (n=6).

**Supplementary Figure 5:**
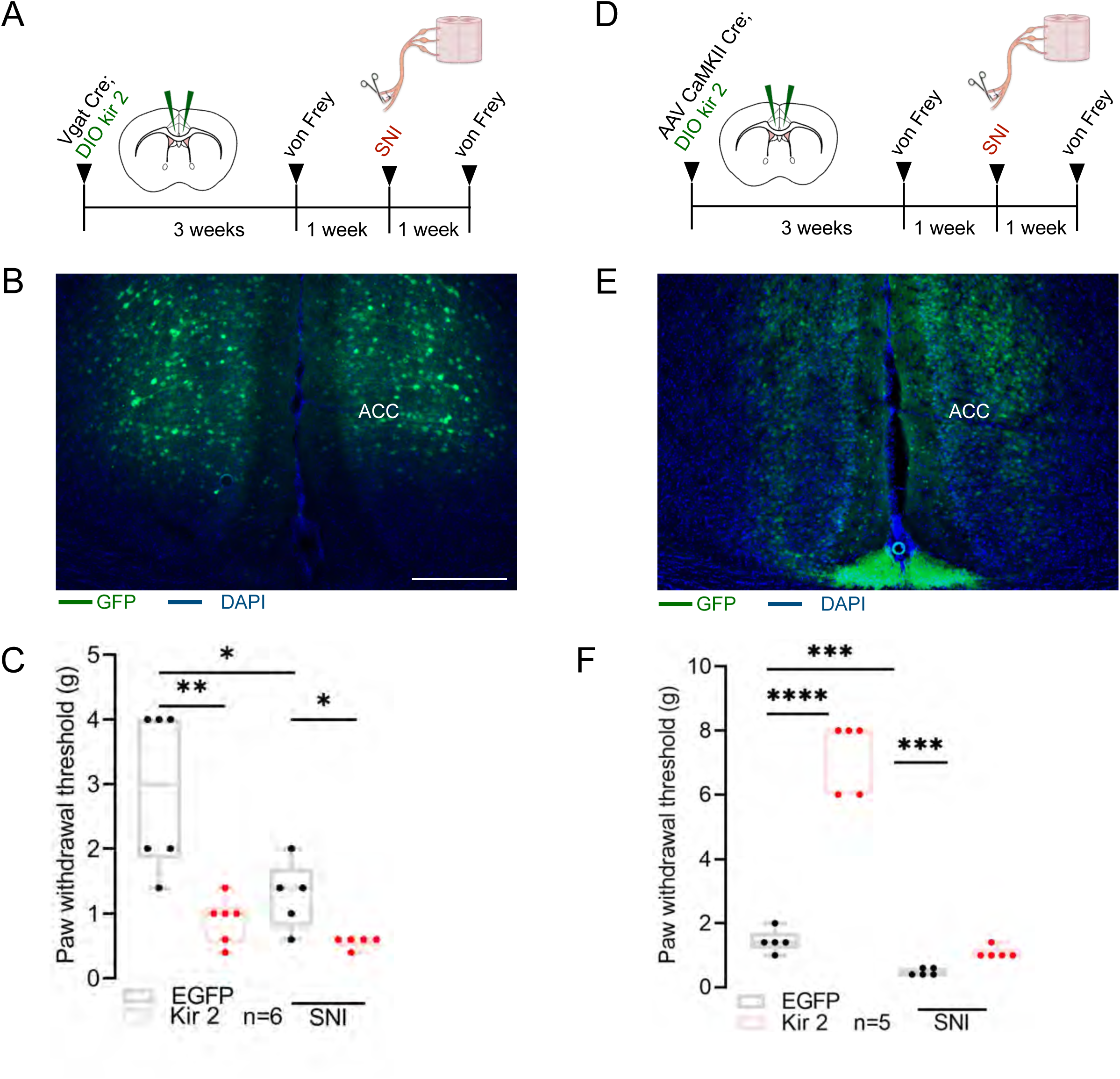
related to- Fig 7. Silencing ACC^D1R^ neurons is sufficient to induce mechanical allodynia. A) Schematic of experimental design. AAV encoding CAMKII Cre and Cre-dependent kir2 was injected bilaterally into the ACC of CD1 mice. B) Fluorescence microscope image of AAV DIO Kir 2 bilateral expression in ACC CAMKII neurons. C) Paw withdrawal threshold following inactivation, pre (unpaired t-test, 5.760± 0.5154, ****p<0.0001) and post (unpaired t-test, 0.6000± 0.09381, ***p<0.0002) SNI (n=5). D) Schematic of experimental design. AAV encoding Cre-dependent kir2 was injected bilaterally into the ACC of Vgat Cre mice. E) Fluorescence microscope image of AAV DIO Kir 2 bilateral expression in ACC Vgat neurons. F) Paw withdrawal threshold following inactivation, pre (Unpaired t-test, −2.000± 0.5203, **p=0.0032, n=6), and post (Unpaired t-test, −0.7200± 0.2366, *p=0.0160, n=5) SNI.

**Supplementary Figure 6:**
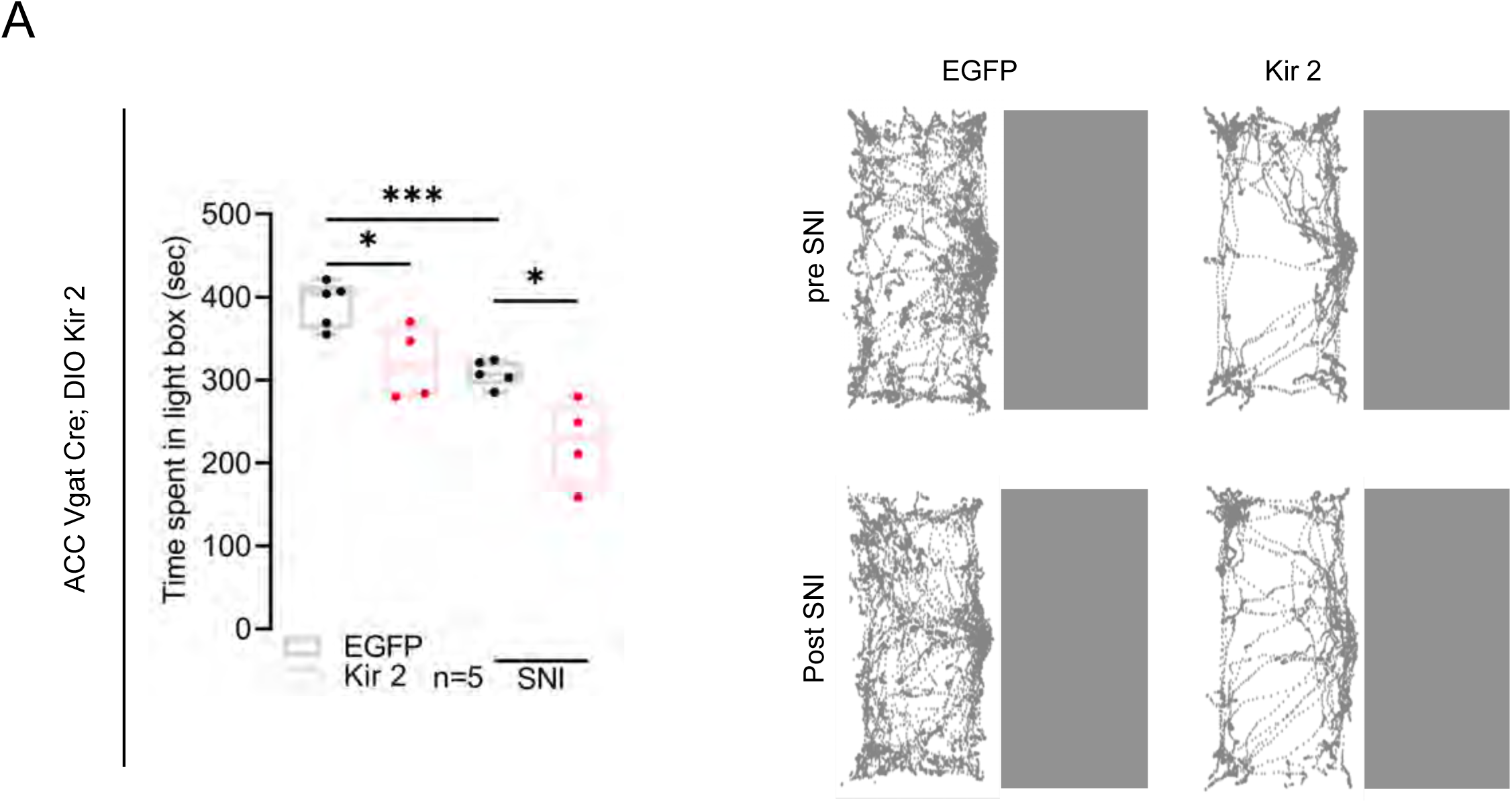
related to- Fig 7. Silencing inhibitory neurons in ACC induces anxiety irrespective of nerve injury. A) AAV encoding Vgat Cre and Cre-dependent excitatory Kir2 was injected bilaterally into the ACC of CD-1 mice. Time spent in the light box before (Unpaired t-test, −70.95± 24.36, *p=0.0226) and after (Unpaired t-test, −83.25± 24.23, *p=0.0109) SNI (EGFP controls, n= 5; Kir2, n=4). On the right side are the trajectory traces of mice navigating through the light-dark arena before and after SNI.

**Supplementary Figure 7:**
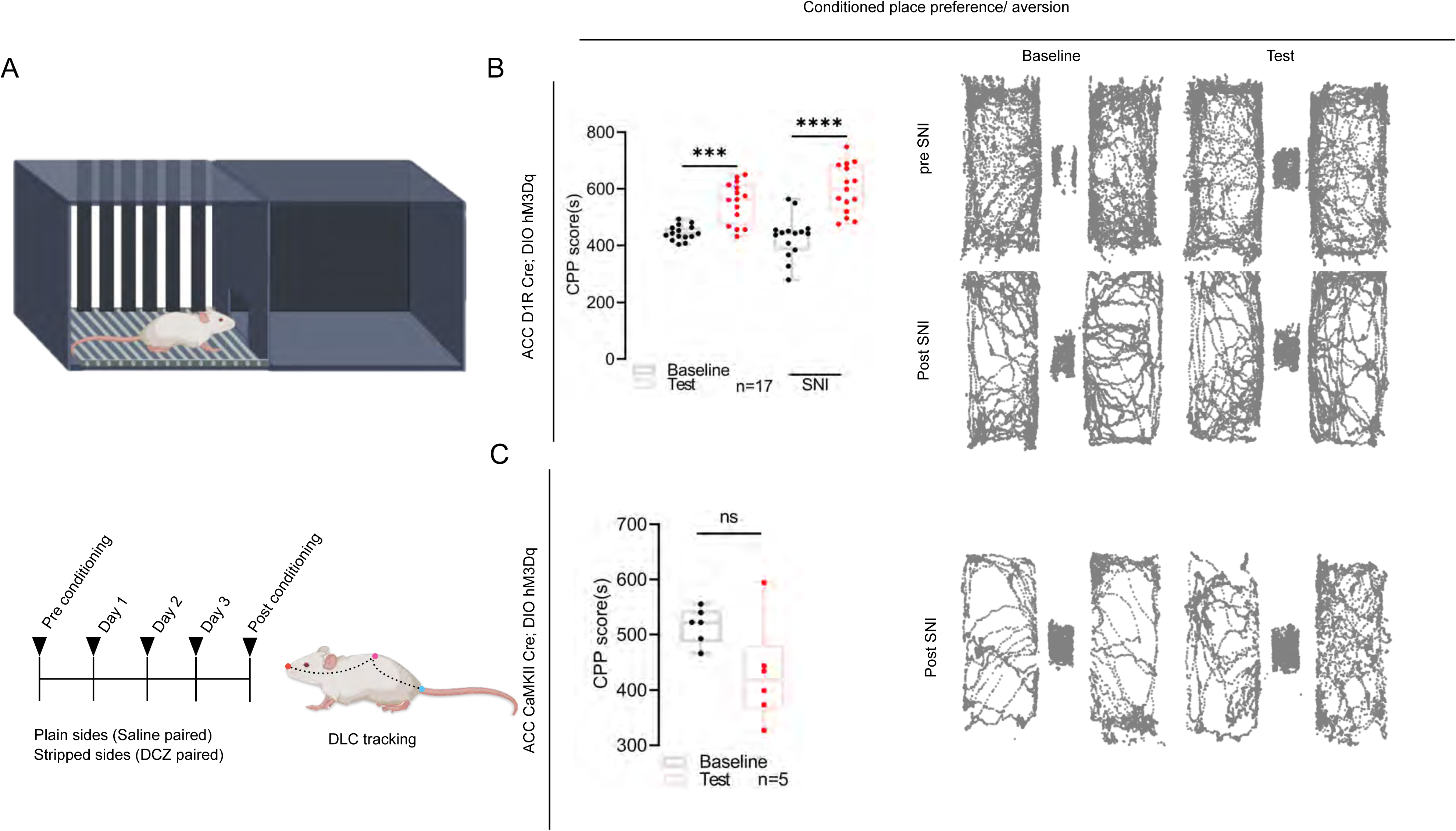
related to- Fig 8. ACC^D1R^ neurons bidirectionally regulate neuropathic pain-induced negative emotions. A) Schematic of CPP experimental design. B) AAVs encoding D1R^Cre^ and Cre-dependent excitatory DREADD hM3Dq were injected bilaterally into the ACC of CD-1 mice. Time spent in the DCZ paired chamber before (Tukey’s multiple comparison, −107.0± 24.95, ***p=0.0004) and after (Tukey’s multiple comparison, −169.2± 24.95, ****p<0.0001) SNI (n= 17). On the right side are the trajectory traces of mice navigating through saline and DCZ paired chambers before and after SNI. C) AAV encoding CAMKII Cre and Cre-dependent hM3Dq-mCherry was injected bilaterally into the ACC of CD-1 mice. Time spent in the DCZ paired chamber after SNI (Paired t-test, −87.57± 35.85, p=0.0585, n= 6). On the right side are the trajectory traces of mice navigating through saline and DCZ paired chambers after SNI.

**Supplementary Figure 8:**
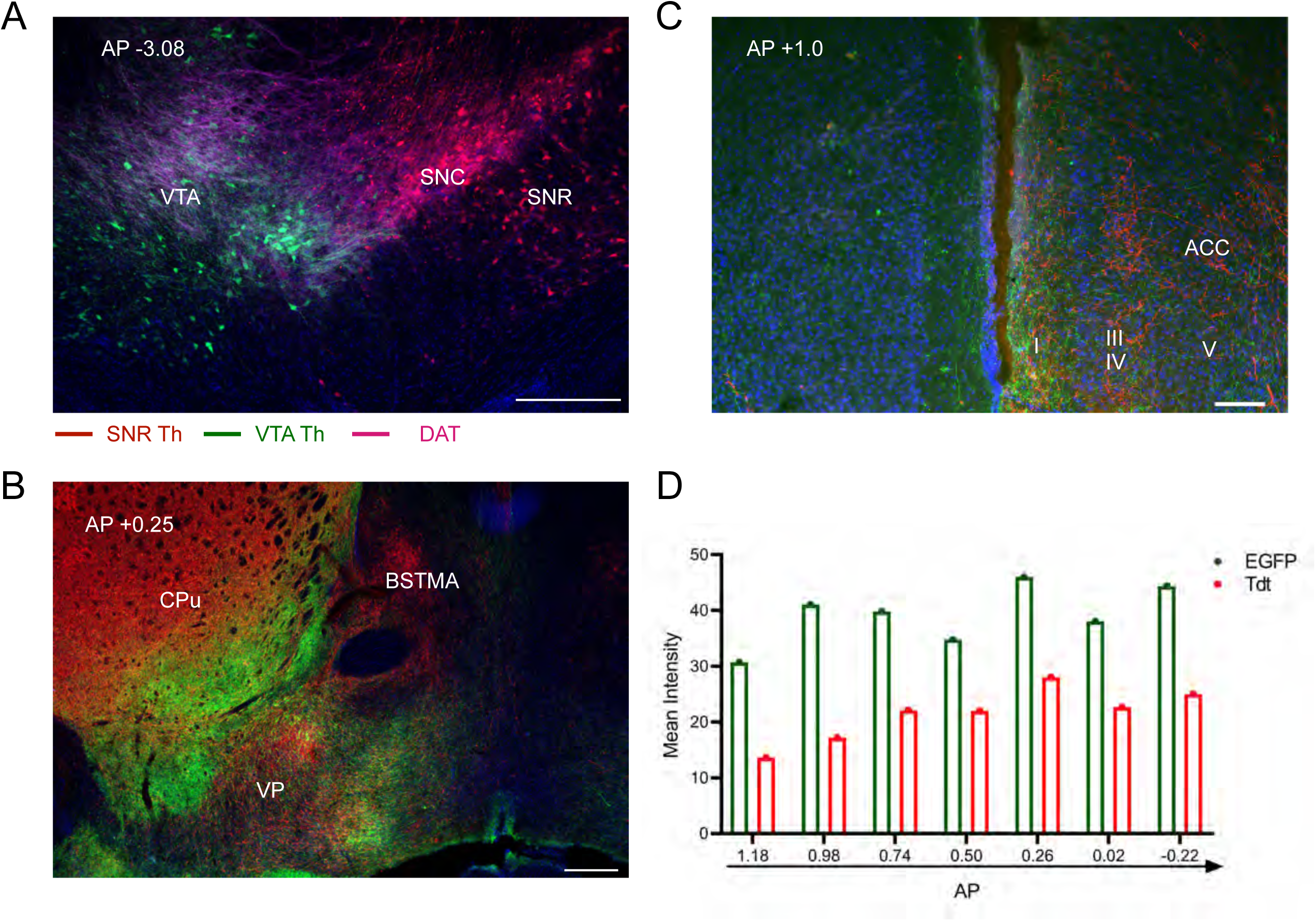
related to Fig 10. Axonal targets of ACC^D1R^ neurons. A) Confocal image showing labelling of Th-positive neurons in VTA (green), SNR, and SNC (red), along with the Dopamine transporter (DAT). B) Fluorescence microscope image showing dense projections of SNR Th neurons into the Caudate putamen (CPu) and BSTMA. C) Fluorescence microscope image showing VTA (green) and SNR (red) Th neurons’ projections in ACC. D) Whole brain quantification of the VTA (green) and SNR (red) Th neurons’ inputs to the ACC. Data are presented as mean ±SEM, from n=2 mice.

